# Shepherding by biased diffusion organizes diffusible proteins on microtubules

**DOI:** 10.1101/2025.06.28.662138

**Authors:** Leila Farhadi, Shane A. Fiorenza, Sithara Wijeratne, Konstantinos Nakos, Yang Yue, Morgan Pimm, Jakia Keya, Kristen Verhey, Radhika Subramanian, Meredith D. Betterton

## Abstract

How microscopic interactions give rise to cellular-scale order in the crowded environment of cells remains an open problem. Far-from-equilibrium mixtures of active and passive molecules self-organize, but the principles remain unclear. Microtubules are cytoskeletal polymers composed of 13 protofilaments bound by mixtures of actively moving motors and passively diffusing microtubule-associated proteins (MAPs), serving as a model system for self-organization on a multi-lane lattice. Here, we combine computational modeling, analytic theory, and *in vitro* reconstitution to demonstrate that motors can “shepherd” diffusive MAPs by rectifying Brownian motion into net directional drift without motor-MAP binding. Our model reveals that biased movement increases when fewer protofilaments are accessible or when lateral diffusion between protofilaments is limited, identifying local dimensionality and diffusion as key parameters governing spatial patterning. Notably, shepherding depends not on individual motor processivity but on the total number of motors bound to microtubules. We experimentally recapitulate micron-scale shepherding with a kinesin-1 motor (K401) and a diffusive MAP (PRC1), which show no detectable binding. These findings reveal shepherding as an emergent mechanism by which ensembles of motors generate micron-scale spatial patterning and transport of MAPs without direct binding interactions.

## INTRODUCTION

Collections of particles interact to generate order and spatial patterns across scales in physics. Understanding the rules by which microscopic interactions generate collective behavior is an important problem for both condensed-matter physics and cell biology. Systems of active (driven or self-propelled) and passive (diffusive) particles can exhibit both activity-driven biased motion and spatial organization of the passive particles^1–3^. However, despite extensive theoretical and computational study of these systems, relatively few experimental realizations robustly show both biased transport and spatial patterning^1,4^, which has limited understanding of the physical principles of active-passive mixtures^5^.

Cells contain many types of molecules, providing new problems in nonequilibrium self-organization. A prototypical example is cytoskeletal microtubules, which are biologically important for cell shape, movement, division, and intracellular transport. Many proteins bind to microtubules to regulate their structure and organization, while motor proteins move along them to transport cargo within the cell. As a result, the spatial distribution of proteins on cytoskeletal microtubules plays an important role in their emergent function. However, the contributions of motor activity, molecular interactions, crowding, and geometry to emergent mesoscale order remain unresolved. Because microtubules have ∼13 parallel protofilaments, they are multi-lane lattices with geometry intermediate between one and two dimensions, depending on how proteins switch between protofilaments. Protein motion and patterning occurs both longitudinally and laterally across protofilaments^6–8^. In such crowded, far-from-equilibrium mixtures of active motors and other bound proteins, the physical control parameters that govern patterning and localization are poorly understood.

A key gap in knowledge is how interactions of active motor proteins with diffusively bound microtubule-associated proteins (MAPs) result in self-organization. Non-motor MAPs are a large class of proteins that bind to microtubules and perform a range of functions such as regulation of microtubule dynamics and mediation of microtubule bundling^9–12^. A characteristic property shared by many non-motor MAPs is their non-directional diffusive movement on microtubules^13–19^. Work on motor-MAP interactions has typically focused on either cargo transport by highly processive motors via direct binding interactions^20–25^, or alternatively, how motors navigate “roadblocks” of MAPs^6,26–28^. However, a few studies suggest that non-directional diffusive movement could aid spatial patterning on MAPs. For example, microtubule depolymerization can induce the relocalization of MAPs because the microtubule tip acts as a diffusion barrier. This plays a role in concentrating MAPs on microtubule ends and in the movement of kinetochore proteins due to microtubule depolymerization^29–34^. A small number of studies have examined other interactions in motor-diffusive MAP mixtures. Imaging of fluorescently tagged kinetochores in *Saccharomyces cerevisiae* lysates suggested that micron-scale movement of individual kinetochore molecules on microtubules may result from transient interactions with a kinesin motor^35^. Further, a mathematical model of a motor and a diffusive MAP bound to a microtubule represented as a single protofilament showed that motor-driven accumulation of the MAP occurs without direct binding interactions^36^. However, conclusive evidence of transport and patterning of diffusive MAPs by motors in the absence of direct binding is currently lacking. While diffusion-based mechanisms are likely to depend strongly on dimensionality, for example through protofilament number, these parameters have not been extensively investigated in the context of molecular patterning on microtubules. As a result, the range of mechanisms driving motor-MAP self-organization is unknown, and the rules governing spatial patterning on microtubules remain unclear.

Here we identify a mechanism for diffusive MAP patterning on microtubules by motors without direct motor-MAP binding. In both a multi-protofilament lattice model and reconstitution experiments, we find that repeated local steric interactions with motors rectify the Brownian motion of diffusive MAPs into net drift aligned with the stepping direction of the motors. This produces MAP streams along the lattice and enhanced concentration of the MAP at the plus end, a process we call “shepherding.” Both MAP directional movement and MAP accumulation at the microtubule end increase when the microtubule becomes more quasi-one-dimensional (due to fewer accessible protofilaments) or lateral diffusion between protofilaments is decreased. These findings reveal a novel mechanism in which directional transport and spatial patterning of microtubule-bound proteins is achievable without direct motor-MAP binding.

## RESULTS

### A computational model predicts motor-driven transport and patterning of MAPs via biased diffusion

Previous theoretical work found that a processive motor can concentrate a diffusive MAP near the end of a single protofilament of α,β-tubulin heterodimers^36^. Because diffusion is sensitive to dimensionality and geometry, we sought to test this mechanism in a multi-protofilament model. We modeled 8 protofilaments to represent a surface-attached microtubule for which protein binding may be inaccessible to about half the protofilaments near the surface. We considered two proteins (Fig. 1): a plus-end-directed motor and a passive MAP that can diffuse on the lattice but does not directly bind to the motor (Supplementary Material, Table S1, parameters chosen based on comparison with experiments as discussed below). In our model, motors move toward the plus-end of the lattice along a single protofilament via stepping of two heads which undergo a mechanochemical ATP hydrolysis cycle, while MAPs diffuse both longitudinally (along a protofilament) and laterally (hopping between protofilaments, Fig. 1A,B)^37^. The motors and MAPs sterically exclude each other on the lattice, but do not otherwise interact. In addition to steric exclusion, bound MAPs have cooperative interactions that increase their microtubule-association lifetime when adjacent to other bound MAPs, consistent with the measurements of a model diffusive MAP, Protein Regulator of Cytokinesis 1 (PRC1)^16^. MAP binding occurs at a constant rate proportional to the solution concentration. Because MAPs have cooperative interactions with neighbors, the unbinding rate depends on the number of bound lattice neighbors (up to 4 including both longitudinal and lateral neighbors, Supplementary Material). Our simulations, carried out in the open-source CyLaKS framework, reproduce the results of previous work^36^ in the appropriate limit (Supplementary Material, Fig. S1A-D, Github: Betterton-Lab/CyLaKS) ^37^. Our simulations showed that over ∼2 min a concentrated region of MAPs forms near the plus end of the microtubule, illustrating spatial patterning in an “endzone” (Fig. 1C)^37^. ^36^This occurs because motors adjacent to MAPs prevent diffusive steps toward the microtubule minus-end (Fig. 1C). When MAPs diffuse towards the plus end, motors can step forward and lock in the displacement of MAPs. Motors therefore bias MAP diffus ion to *shepherd* MAPs along the microtubule. In addition, while MAP cooperativity is not strictly required for endzone formation, it does significantly enhance endzones for higher MAP concentration (Fig. S1E,F).

**Fig. 1.**
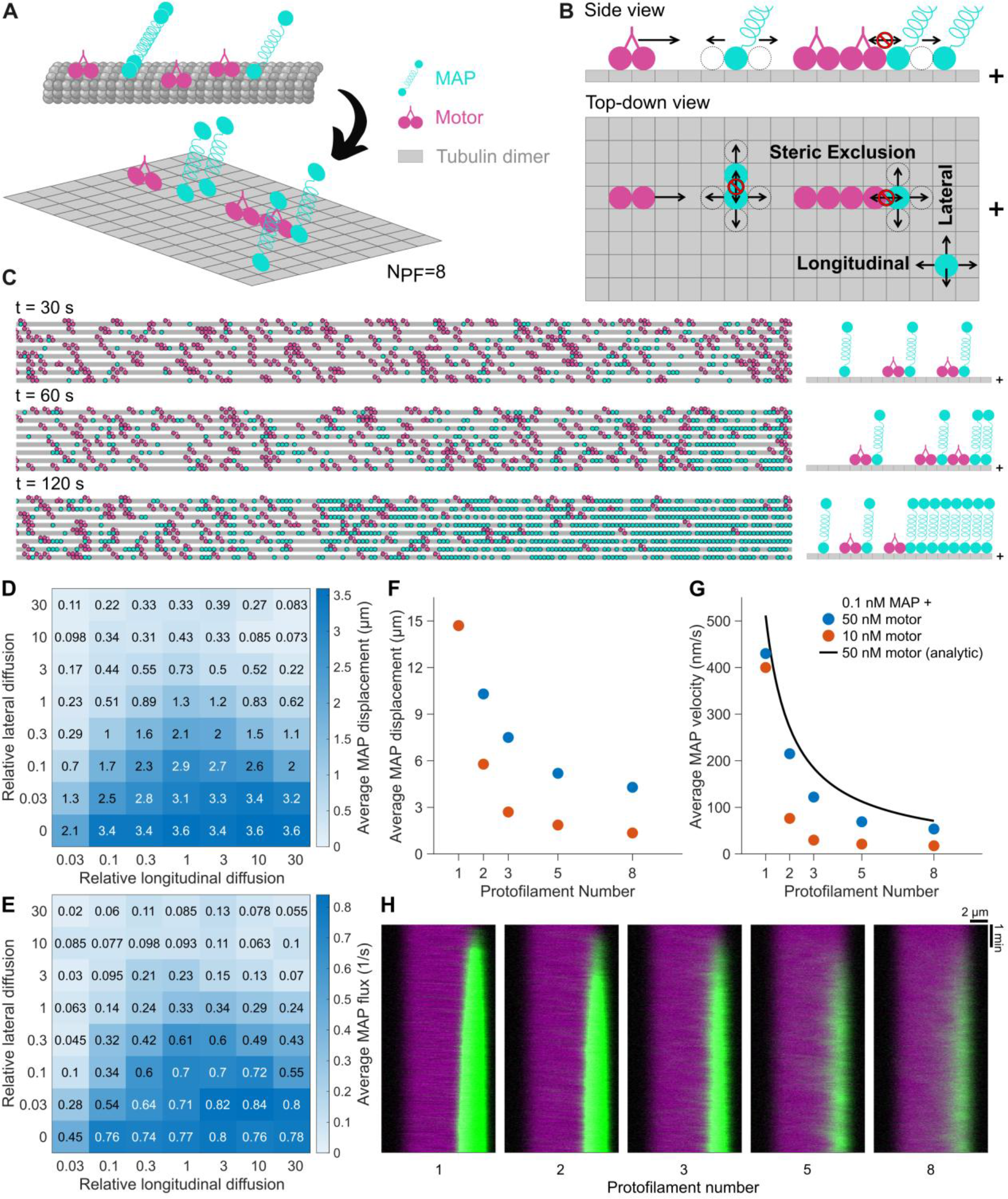
A multi-protofilament biophysical model shows that motors can bias the diffusion of MAPs for their accumulation into an endzone. (A-B) Model schematic. The surface-attached microtubule is modeled with 8 accessible protofilaments. Each lattice site represents a tubulin heterodimer. Motors (magenta) step toward the plus end of the microtubule (right pointing arrows). MAPs (cyan) diffuse both longitudinally (along a protofilament, left and right pointing arrows) and laterally (between protofilaments, up and down pointing arrows in top-down view). The motor and MAP interact only by steric exclusion that prevents them from occupying the same lattice site (disallowed displacements marked with crossed-out circles). (C) Snapshots of simulation (left) and schematic (right) showing that shepherding over time concentrates MAPs near the plus end of the microtubule. MAP and motor concentration, 0.1 nM and 100 nM. (D, E) Average MAP displacement (D) and flux on protofilament (E) as a function of relative lateral and longitudinal diffusion coefficient for microtubules 1000 sites≈8 μm long. Motor concentration, 10 nM. MAP concentration, 0.1 nM. (F, G) Average MAP displacement (F) and velocity (G) as a function of number of protofilaments for 0.1 nM MAP and either 10 nM (orange) or 50 nM (blue) motor concentration for microtubules that are 5000 sites≈40 μm long. (H) Simulated kymographs of MAP accumulation at endzones over time for simulations with varying protofilament number. MAP (green) and motor (purple) concentration, 0.1 and 50 nM. Microtubules are 1000 sites≈8 μm long.

Our proposed shepherding mechanism depends on how effectively motors can bias MAP diffusion. To explore this, we varied the longitudinal and lateral diffusion coefficient of MAPs, starting from parameters for which these coefficients are equal (*D*_long_ = *D*_lat_, Table S1, Supplemental Material). To quantify transport by shepherding, we measured both the average MAP displacement toward the microtubule plus end before unbinding and the average MAP flux across the microtubule midzone. These quantities are zero for unbiased diffusion and increase for stronger shepherding. In our model, shepherding occurs over a wide range of diffusion coefficients but is enhanced for lower lateral diffusion. Increasing lateral diffusion tended to decrease shepherding displacement and flux, because MAPs hopped laterally away from the protofilament-tracking motors (Fig. 1D,E). We also quantified MAP lifetime and shepherding velocity as a function of the diffusion coefficient (Fig. S2A-C). Notably, both the displacement and flux were non-monotonic in the longitudinal diffusion coefficient over a range of lateral diffusion coefficient. If the longitudinal diffusion coefficient was too low, MAPs did not hop fast enough for significant transport. However, if the longitudinal diffusion coefficient was too high, they quickly diffused away after lateral hops, negating any prior transport. Thus, an intermediate “Goldilocks” diffusion coefficient allows for the most efficient transport via shepherding. The previously measured PRC1 diffusion coefficient is in this optimal range^38^.

Due to similar trends in displacement and flux, we choose MAP displacement as a primary measure of shepherding transport in this work. We also note that higher shepherding displacement typically leads to greater accumulation of MAPs at the endzone of microtubules (Fig. S2D), showing that transport and endzone accumulation are correlated. Through comparison of this endzone accumulation to our experimental measurements (below), we found a lateral diffusion coefficient five times higher than the longitudinal one provided the best fit (Table S1) and use this value in what follows. This parameter choice leads to relatively weak but distinctly observable shepherding at MAP and motor concentrations of 0.1 and 10 nM, respectively. Decreasing this ratio, i.e., making the lateral and longitudinal diffusion coefficients more similar, would lead to stronger shepherding across all conditions. We note, however, that the exact value of the lateral diffusion coefficient chosen will not significantly change the conclusions presented. Given the dependence of shepherding on the diffusion parameters of MAPs, we asked whether this mechanism could drive sorting of different types of MAPs. In simulations with two MAPs with different lateral and longitudinal diffusion coefficients, shepherding can cause sorting of the MAPs into two distinct regions in the endzone (Fig. S2E).

The importance of the balance between lateral and longitudinal diffusion suggested that the effective dimensionality via the number of locally accessible protofilaments could control shepherding. For example, while we assume that the surface-attached microtubules studied in our experiments have roughly half of their protofilaments accessible, only 1 or 2 might be accessible in dense microtubule bundles, potentially leading to different shepherding behavior based on local geometry. To explore this, we formulated an analytic model to estimate the shepherding velocity and its dependence on protofilament number (Supplementary Material) and ran simulations with varying protofilament number for two sets of MAP and motor concentrations that correspond to weak and intermediate shepherding conditions (Fig. 1F-H). Due to the significantly enhanced shepherding for single protofilaments, longer microtubules of 5000 sites (≈40 *μ*m) were used. We found that shepherding transport and velocity decrease as the number of protofilaments in our model increased, consistent with our results on the MAP diffusion coefficient. The analytic model shows similar qualitative trends. Notably, shepherding on the micron scale can still be observed at the highest protofilament number tested. Therefore, shepherding strongly depends on the number of accessible protofilaments, facilitating different patterning based on local geometry.

### Shepherding depends on the total number of bound motors, not motor processivity

To further explore the principles of transport and patterning in shepherding, we varied MAP and motor parameters in our simulation. First, we varied MAP bulk concentration and lifetime (Fig. 2A,B). Higher MAP lifetime favored shepherding since binding events were longer, allowing larger displacement. However, appreciable transport can still be achieved for lower MAP lifetime. Bulk MAP concentration in the 10 pM-1 nM range had little effect on shepherding, except at high lifetime and concentration where the lattice became crowded and shepherding was ineffective due to a nearly saturated lattice (Fig. S3A,B). Measured MAP displacement was further limited by the formation of an extended endzone, which restricted how far MAPs could travel before becoming immobilized due to steric effects (Fig. S3C). Shepherding is therefore possible across a wide range of MAP lifetimes and bulk concentrations.

**Fig. 2.**
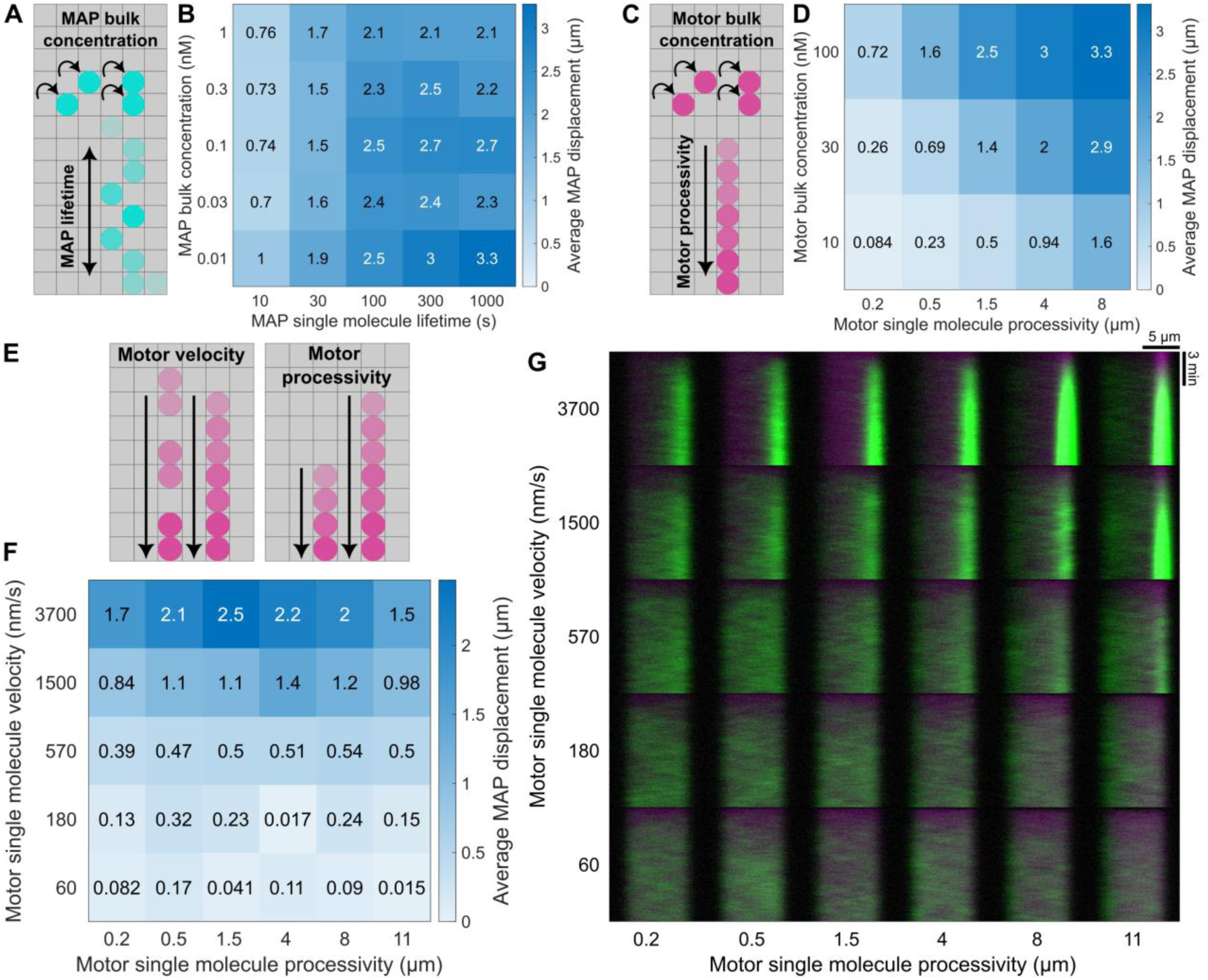
Shepherding efficiency is set by number of motors bound to microtubule, not individual processivity. (A) Schematic of MAP bulk concentration and lifetime parameters. (B) Average MAP displacement as a function of MAP concentration and lifetime. Motor concentration, 100 nM. Note that apparent differences in shepherding result from variability in the stochastic model. The standard error of the mean is larger for 0.01 nM MAP concentration due to the smaller sample size. (C) Schematic of motor concentration and processivity parameters. (D) Average MAP displacement as a function of motor concentration and lifetime. MAP concentration, 0.1 nM. (E) Schematic of motor velocity and processivity parameters. (F) Average MAP displacement as a function of motor velocity and processivity. (G) Simulated kymographs of MAP accumulation over time for simulations with varying motor velocity and procssivity. MAP (green) concentration, 0.1 nM. For motor (purple) concentrations see Fig. S3 caption. For all simulations, microtubules are 1000 sites≈8 μm long.

We next varied motor bulk concentration and processivity to see their contribution to shepherding (Fig. 2C,D). Motor stepping is essential for shepherding because directional movement of motors breaks the left-right symmetry along the filament which is required to bias MAP diffusion. Highly processive motors meet this requirement, but our model suggests that motors of lower processivity can also drive shepherding if present in sufficient numbers to provide a mobile diffusion barrier. If motor processivity and bulk concentration were both low, shepherding did not occur due to a lack of sufficient motors to bias MAP diffusion. If motor concentration and processivity were high enough to favor shepherding, a decrease in motor processivity could be compensated for by increased bulk concentration. Notably, shepherding transport strongly correlated with the number of motors bound to the microtubule rather than the single-motor processivity (Fig. S3D-F). In fact, for strong shepherding, measured motor processivity significantly decreased due to crowding effects (Fig. S3F,G). Strong shepherding despite decreased processivity is possible because motors can “pass off” MAPs to one another, allowing one motor to continue shepherding the MAP after the first unbinds from the microtubule. Therefore, robust MAP transport via shepherding is achievable even for motors with low processivity.

To investigate how motor properties affect shepherding independent of the number of bound motors, we varied motor velocity and processivity while keeping the total number of motors bound approximately constant (Fig. 2E-G). The number of bound motors along the 8-micron-long microtubule was kept constant at approximately 400 for all simulations (Fig. S3H,I) by adjusting bulk motor concentration based on the measured lifetime of motors, which depends on both processivity and velocity (Fig. S3J, Supplemental Materials). As expected, motor velocity strongly tuned shepherding, because faster movement more strongly biased MAP diffusion. Consistent with our results above, motor processivity only had a modest effect on shepherding displacement (Fig. 2F) when the total number of motors bound to the lattice was kept constant. Remarkably, when motor processivity was decreased to just 200 nm, strong shepherding still occurred when motor occupancy was maintained. For high processivity, we found enhanced endzone formation (Fig. 2G) that resulted in slightly decreased measured MAP displacement due to crowding effects limiting motion. While conventional transport can occur with a single high-processivity motor bound to a cargo, shepherding depends on multiple motors bound to the lattice and is more sensitive to the total number of bound motors rather than single-motor processivity. For example, a motor with modest processivity (200 nm) can transport MAPs via shepherding for motor density corresponding to just 5% average occupancy with 2.5% MAP occupancy (Fig. S3E). Shepherding is a collective effect of multiple motors, not a result of a single motor transporting a bound MAP.

### Translocation of PRC1 to microtubule plus-ends occurs in the absence of canonical motor-cargo interactions

To experimentally test the shepherding mechanism suggested by our model, we designed *in vitro* reconstitution assays with PRC1 and a minimal dimeric kinesin-1 recombinant protein, which includes only the motor domain and the first coiled-coil dimerization domain of Drosophila kinesin-1 (hereafter referred to as K401, Fig. 3, S4A-E). This truncated kinesin-1 construct was chosen to minimize the possibility of binding interactions between kinesin and PRC1. To examine the localization of PRC1 on microtubules in the presence of K401, we immobilized GMPCPP-polymerized taxol-stabilized microtubules, that were labeled with X-rhodamine and biotin, on a neutravidin-coated glass coverslip (Supplementary Methods). Then, an assay buffer containing a mixture of GFP-PRC1, K401-Clip-647 (K401-Clip protein labeled with CLIP-Surface 647), and 1 mM ATP was flowed into the imaging chamber and imaged using Total Internal Reflection Fluorescence Microscopy (TIRFM, Fig. 3A).

**Fig. 3.**
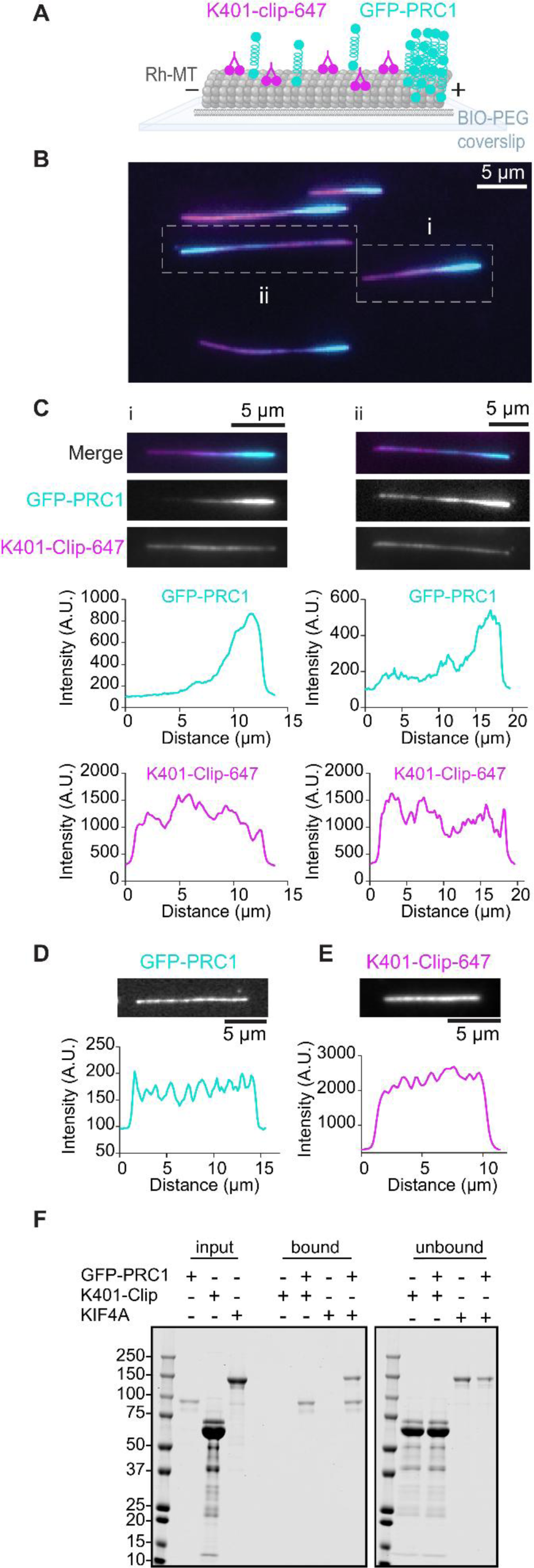
Translocation of PRC1 to microtubule plus-ends in the absence of canonical motor-cargo interactions. (A) Experimental set up for visualizing GFP-PRC1 (cyan) shepherding by K401-Clip-647 (magenta). GMPCPP-polymerized and taxol-stabilized biotinylated microtubules are immobilized on a Biotin-PEG coated coverslip using neutravidin. Scale bars are 5 μm. (B) A representative field of view from the shepherding assay showing microtubule-associated GFP-PRC1 (cyan, 1 nM) and K401-Clip-647 (magenta, 100 nM). (C) Two examples from the image in 1B highlighted along with individual GFP-PRC1 and K401-Clip-647 images in greyscale. Corresponding intensity versus distance along the microtubule plots are shown. (D-E) Representative microtubule from control experiments with (D) 1 nM GFP-PRC1 and (E) 100 nM K401-Clip-647. (G) Biochemical pull-down assay for GFP-PRC1 and K401-Clip. GFP-PRC1 (150 nM) was immobilized on GFP-trap agarose beads and subsequently combined with 5 µM K401-Clip or 0.5 μM KIF4A, a known PRC1 binding protein, in the absence of microtubules. Coomassie-stained PAGE gel of input, bound, and unbound fractions from the pull-down assay are shown.

When immobilized microtubules were incubated with a solution containing 100 nM K401-Clip-647 (magenta), 1 nM GFP-PRC1 (cyan) and 1 mM ATP, we observed a striking accumulation of PRC1 molecules at microtubule ends over time (Fig. 3B,C). Protein concentrations were chosen to optimize spatial patterning. In contrast, the distribution of K401 motors remained uniform over the entire microtubule length. Control experiments with only one protein (GFP-PRC1 or K401-Clip-647, Fig. 3D,E) revealed uniform protein binding along the microtubule. To confirm that PRC1 accumulates at the plus-ends of microtubules, we performed a microtubule gliding assay. In this assay biotinylated K401 molecules were immobilized on the glass coverslip using a neutravidin linkage, followed by addition of non-biotinylated microtubules, GFP-PRC1 and K401. Time-lapse imaging showed that the accumulated PRC1 (cyan) was at the rear end of gliding microtubules, which is the plus-end for K401-based motility (Movie S1).

The minimal K401 dimer is not expected to bind PRC1. Consistent with this, no enrichment of K401 was seen along with PRC1 at microtubule plus ends (Fig. 3C). To explore the possibility of PRC1-K401 interaction more rigorously, we performed biochemical pull-down assays (Fig. 3F). GFP-PRC1 (150 nM) was immobilized on GFP-trap agarose beads and subsequently combined with 5 µM K401-Clip or 0.5 μM KIF4A, a known PRC1 binding protein, in the absence of microtubules. No detectable binding was observed between 150 nM GFP-PRC1 and 5 µM K401-Clip. Control experiments with 150 nM GFP-PRC1 and 0.5 μM KIF4A showed a direct interaction between the two proteins as expected^24^ .

To observe the dynamics of K401-mediated PRC1 transport and patterning in real time, we acquired time lapse images immediately after the addition of proteins (100 nM K401-Clip-647 and 1 nM GFP-PRC1) and ATP to immobilized microtubules (Fig. S5). We observed the appearance of the PRC1 endzone at one end of each microtubule within the first 30 seconds of protein addition (Fig. S5A,B). The length and intensity of the PRC1 endzone increased initially, then reached steady state. We quantitatively examined changes in overall protein occupancy on the microtubule with time (Fig. S5C,D). We observed that the PRC1 fluorescence density (intensity per unit length) on microtubules increased with time (Fig. S5C,I). Control experiments with 1 nM GFP-PRC1 alone showed a similar trend (Fig. S5K). In contrast, K401-Clip-647 microtubule occupancy showed a slight decrease with time (Fig. S5D,J). Control experiments with 100 nM K401-Clip-647 alone indicate that this reduction may arise from photobleaching (Fig. S5L). The trends in occupancy over time were similar in our computational model (Fig. S5E-H).

Collectively, these data show that PRC1 can be transported and concentrated in endzones by K401, despite the lack of detectable K401-PRC1 binding in solution. These observations provide experimental evidence for shepherding-based MAP transport and patterning.

### Shepherding occurs because motors bias the diffusion of PRC1

In our computational model, shepherding occurs by biased diffusion (Fig. 1A-C). We therefore imaged PRC1 movement on microtubules to test whether single molecules moved by biased diffusion. For this experiment, we focused on three experimental conditions for which endzone formation was relatively slow (Fig. 4). To resolve the movement of single PRC1 molecules, we used speckle imaging by mixing low levels of fluorescent Snap-647-PRC1 with non-fluorescent PRC1 such that 10-34% of PRC1 molecules were labeled. To capture PRC1 movement during endzone formation, we imaged for 5 min immediately after addition of PRC1 and non-fluorescent K401 (Fig. 4A-C, S6A, and Movie S2, S3). Snap-647-PRC1 molecules moved and accumulated at one end of the microtubules. Kymographs from these experiments showed directed streams consistent with biased diffusion of PRC1 over microns. Such directed tracks were not observed in control experiments with no K401 (Fig. 4D,E). We quantified the average velocity of PRC1 from our kymographs (Supplementary Methods). The PRC1 velocity (∼70-86 nm/s) was not strongly dependent on protein concentration in the measured range (Fig. 4F-H, S6B). Automated particle tracking for PRC1 molecules was challenging because PRC1 tracks frequently crossed during their long-lifetime binding events. However, we were able to analyze short trajectories without crossing events and measured the mean and mean-squared displacement of single PRC1 molecules over seconds to minutes (Fig. S7; Supplementary Methods). In the absence of motors, PRC1 trajectories showed a Gaussian distribution of displacements with zero mean, as expected for unbiased diffusion (Fig. S7A-C). In the presence of motors, the mean of the Gaussian shifted toward positive values and increased over time. Mean-squared displacement was increased in the presence of motors (Fig. S7D). This adds further evidence of biased diffusion due to motors.

**Fig. 4.**
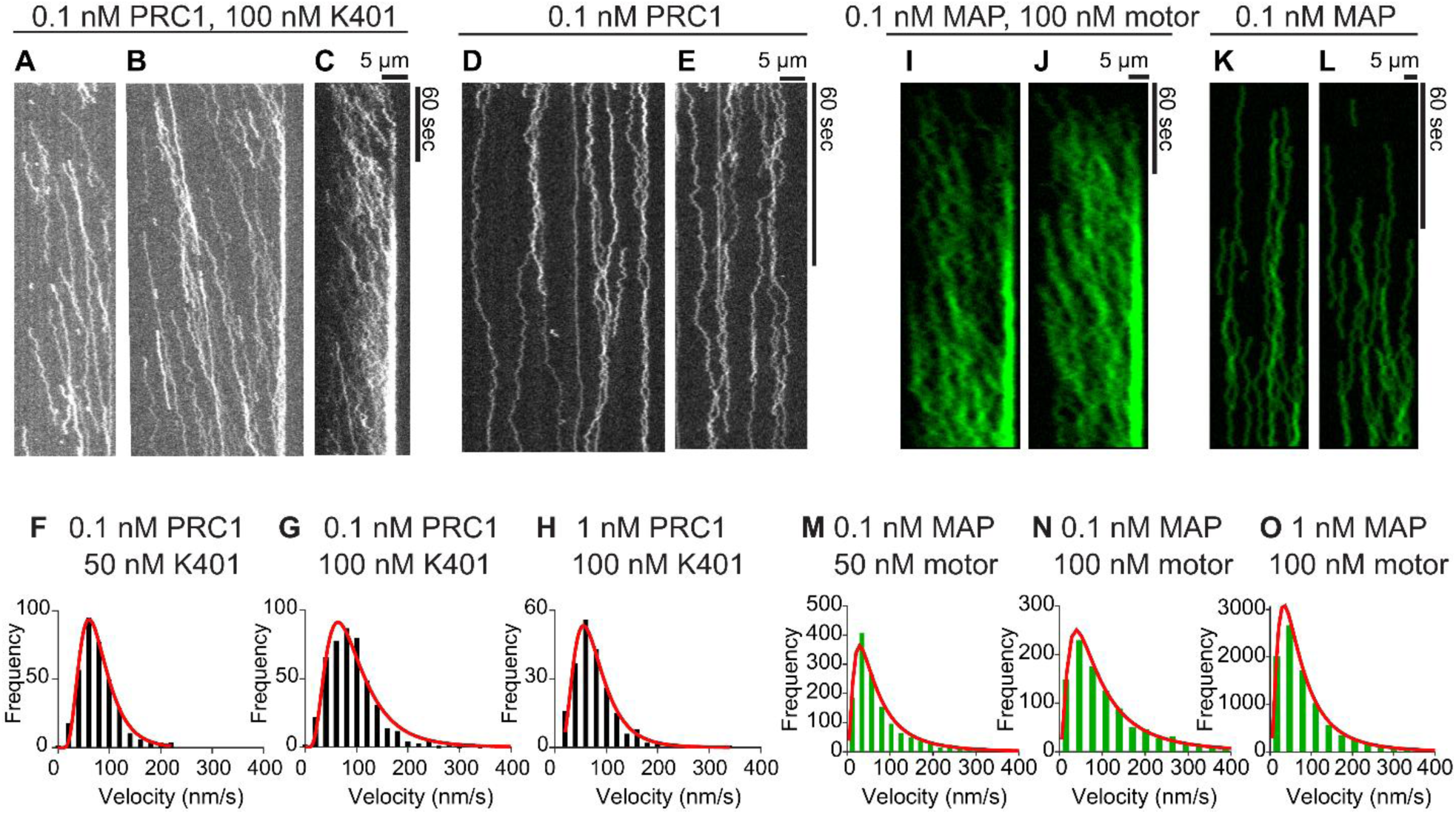
Shepherding occurs by K401-mediated biased diffusion of PRC1. (A-C) Representative kymographs from speckle imaging of fluorescent Snap-647-PRC1 showing biased diffusion of PRC1 (0.1 nM) in the presence of non-fluorescent K401 (100 nM). The fraction of fluorescent PRC1 is 10% in panels A and B, and 34% in panel C. The horizontal and vertical scale bars are 5 μm and 60 s. (D-E) Kymographs of PRC1 molecules in control experiments with 0.1 nM PRC1. The fraction of fluorescent PRC1 is 1.4%. (F-H) Histograms of PRC1 velocity at indicated concentrations. The histograms were fitted to a lognormal distribution (red). Geometric mean, geometric standard deviation and N are 0.1 nM PRC1 and 50 nM K401: 73.6 nm/s, 1.6, N=357; 0.1 nM PRC1 and 100 nM K401: 86.4 nm/s, 1.7, N=466; 1 nM PRC1 and 100nM K401: 71.6 nm/s, 1.7, N=212. N is the number of tracks analyzed in four independent experiments in each condition. The bin size is 20. (I, J) Simulated kymographs from the computational model of MAP molecules showing biased diffusion in the presence of non-fluorescent motors. To match experimental speckle imaging, only 10% of MAP molecules are visible in simulated kymographs. Microtubules in simulations are 4000 sites≈32 μm long. (K, L) Simulated kymographs from the computational model of MAP molecules showing unbiased diffusion in the absence of motors. To match experimental speckle imaging, only 1% of MAP molecules are visible in simulated kymographs. Microtubules in simulations are 3000 sites≈24 μm long. (M-O) Histograms of MAP molecule velocity in the computational model at the indicated concentrations. To mitigate effects from crowding and reflect experimental resolution limitations, only tracks that terminated greater than 3 μm away from the plus end and had an average velocity greater than 15 nm/s are included. The histograms were fitted to a lognormal distribution (red). Geometric mean, geometric standard deviation and N are 0.1 nM MAP and 50 nM motor: 59.8 nm/s, 2.4, N=1419; 0.1 nM MAP and 100 nM motor: 83.2 nm/s, 2.4, N=1004; 1 nM MAP and 100nM motor: 65.0 nm/s, 2.3, N=9344. N is the number of tracks analyzed in simulations of each condition. The bin size is 20 for panel M and 30 for panels N and O. In all cases, data wer taken from 6 independent simulations using different seeds. Microtubules in simulations are 1000 sites≈8 μm long.

To compare the experimental data with predictions of our model, we simulated kymographs of MAP movement with and without motors, for the same MAP and motor concentrations used experimentally. Similar to our experimental observations, motors drove biased diffusion that created directed MAP movement toward plus ends (Fig. 4I,J). Only unbiased diffusion of MAPs occurred in the absence of motors (Fig. 4K,L). We found an average geometric velocity of MAP movement during directed events of 60-84 nm/s, similar to that found experimentally (Fig. 4M-O).

We next experimentally examined how the movement of PRC1 molecules changed once the endzone reached steady state. To avoid artifacts due to photobleaching, different fields of view in the same imaging chamber were imaged at different times (Fig. S6C-E, representative images correspond to different microtubules). In experiments, robust directional transport of Snap-647-PRC1 occurred in the first 5 min. Over time, directed movement slowed or stopped entirely (Fig. S6C). The transition to slower/no movement occurred more quickly, within minutes, for higher PRC1 concentration (Fig. S6E).

These findings suggest that shepherding occurs by K401-mediated biased diffusion of PRC1. Directed movement of PRC1 was most visible early in our experiments and stopped over time, likely due to the increased lattice occupancy of PRC1 relative to K401 (Fig. S6C-E) that limits transport.

### PRC1 shepherding by K401 does not require high motor processivity

Modeling suggested that micron-scale shepherding transport does not require motor processivity on the same length scale (Fig. 2E,F). To compare the movement of PRC1 and K401, we imaged motor activity during shepherding. Experiments were performed with K401-Clip-647 and non-fluorescent K401 such that 0.08-0.3% of motors were fluorescently labeled. For 10 nM K401, we observed similarly processive motor runs with and without 0.1 nM non-fluorescent PRC1, as expected for a processive kinesin (Fig. 5A,B, S8A). However, increasing the K401 concentration to 50 nM led to shorter tracks, which were similar for non-fluorescent PRC1 concentrations ranging from 0 to 1 nM (Fig. 5C-E, S8A). This is consistent with a prior report that K401 processivity is reduced when the lattice is crowded by microtubule-bound proteins^39^. The motor lifetime was ∼1.5 s for 10 nM K401 and decreased to ∼1 s for higher K401 concentration, independent of PRC1 concentration, indicating that K401 lifetime is less sensitive to crowding by PRC1 than K401 crowding (Fig. 5F). Notably, the ∼1 s K401 lifetime is ∼2 orders of magnitude shorter than the PRC1 lifetime, which is of order minutes but is so long that it is difficult to quantify in our experiments (Fig. 4A-E)^16,25^ .

**Fig. 5.**
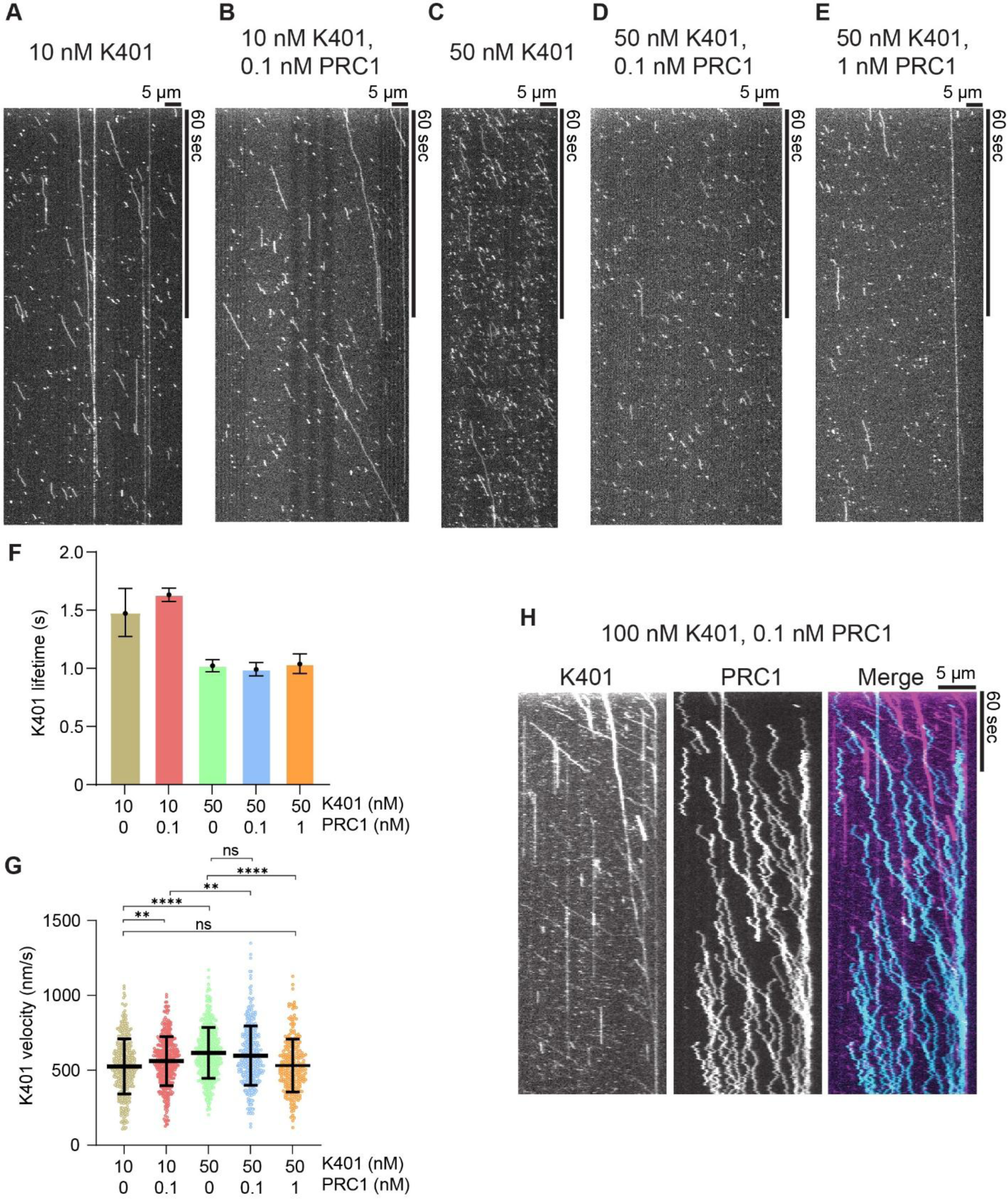
PRC1 shepherding by K401 does not require high motor processivity. (A-E) Representitive kymographs of K401 movement along microtubules at 10 nM K401 (A), 10 nM K401 and 0.1 nM PRC1 (B), 50 nM K401 (C), 50 nM K401 and 0.1 nM PRC1 (D), and 50 nM K401 and 1 nM PRC1 (E). The fraction of fluorescent K401 is 0.08-0.3%. The horizontal and vertical scale bars are 5 μm and 60 s. (F) K401 lifetime versus K401 and PRC1 concentration. Error bars represent 95% confidence interval from exponential fitting to the lifetime histogram. Lifetimes are 10 nM K401: 1.47 s [1.27, 1.69] N=380; 10 nM K401 and 0.1 nM PRC1: 1.63 s [1.58, 1.69] N=398; 50 nM K401: 1.02 s [0.97, 1.08] N=524; 50 nM K401 and 0.1 nM PRC1: 0.99 s [0.94, 1.05] N=387; 50 nM K401 and 1nM PRC1: 1.04 s [0.96, 1.13] N=268. N is the number of K401 tracks analyzed in three independent experiments in each condition. (G) K401 velocity versus K401 and PRC1 concentration. Bars represent mean and standard deviation from fitting to a normal distribution. 10 nM K401: 527 ± 183 nm/s N=380; 10 nM K401 and 0.1 nM PRC1: 568 ± 156 nm/s N=398; 50 nM K401: 606 ± 172 nm/s N=524; 50 nM K401 and 0.1 nM PRC1: 570 ± 184 nm/s N=387; 50 nM K401 and 1nM PRC1: 507 ± 166 nm/s N=268. N is the total number of K401 tracks analyzed in three independent experiments in each condition. P-values were calculated by unpaired parametric t-test. (H) Representative kymographs from speckle imaging of fluorescent K401-Clip-488 and Snap-647-PRC1 showing movement of K401 and biased diffusion of PRC1 molecules at 100 nM K401 and 0.1 nM PRC1. The fraction of fluorescent K401 and PRC1 are 0.03% and 1.9%. The horizontal and vertical scale bars are 5 μm and 60 s.

Next, we sought to determine whether the velocity of directed movement events of K401 differs from that of PRC1. The K401 velocity showed only small changes for the range of concentrations tested (Fig. 5G, S8B). However, K401 motor velocities were ∼500-600 nm/s, ∼8 times faster than the velocity of PRC1 transport under comparable conditions. Therefore, in our experiments, K401 and PRC1 showed both transport velocity and lifetime along the microtubule that differ by a factor of ∼8 and ∼100 respectively. Similarly, in simulations, motor lifetime was 0.7-2.6 s, depending on lattice crowding, and velocity was 530-580 nm/s, while for MAPs lifetime was 80-120 s and velocity 60-85 nm/s. In the model, the significant difference between MAP and motor velocity requires the high lateral/longitudinal diffusion coefficient ratio of ∼5 that we found gave the best model-data comparison. If the lateral and longitudinal diffusion coefficients are comparable, shepherding is enhanced (Fig. S2C), leading to larger MAP velocity in strong shepherding regimes than observed experimentally. To further examine the differences in K401 and PRC1 movement, we simultaneously imaged the two proteins on microtubules using three-color TIRFM (Fig. 5H). Low amounts of K401-Clip-488 and Snap-647-PRC1 were spiked in with non-fluorescent K401 and PRC1 to visualize the movement of individual molecules. As seen in the representative kymographs, the K401 and PRC1 tracks differ in both slope and lifetime, consistent with experiments in which only one of the two proteins was imaged.

Collectively, these findings support the biased-diffusion transport mechanism of shepherding that we propose. The results highlight the differences with conventional motor-cargo transport, which occurs with a similar velocity and lifetime for the bound motor and cargo.

### The relative concentration of K401 and PRC1 determines shepherding efficiency

To further explore how motor and MAP concentration affect shepherding, we examined steady state endzones in experiments with a fixed concentration of GFP-PRC1 and varying amounts of K401-clip-647 (Fig. 6A, S9A). Visually discernible endzones were established under all conditions except at the lowest K401 concentration shown. We performed line scan analysis and determined the percentages of microtubules with quantifiable endzones (Supplementary Methods, Fig. 6B). For 1 nM PRC1 and 1-10 nM K401 (Fig. 6B), only about half of microtubules had endzones (Supplementary Methods), while this increased to >95% for K401 concentration of 100-500 nM (Fig. 6B). While endzones form at 1000 nM K401, they are broad and difficult to clearly identify and quantify for a large fraction of microtubules, leading to an apparent drop in the fraction of microtubules with endzones (Supplementary Methods). At lower PRC1 concentration (0.1 nM), >95% of microtubules had endzones even at 10 nM K401 (Fig. 6B). These findings indicate that efficient shepherding required at least ∼10x greater solution concentration of K401 relative to PRC1 to facilitate biased diffusion of PRC1 (Fig. 6B, S9B).

**Fig. 6.**
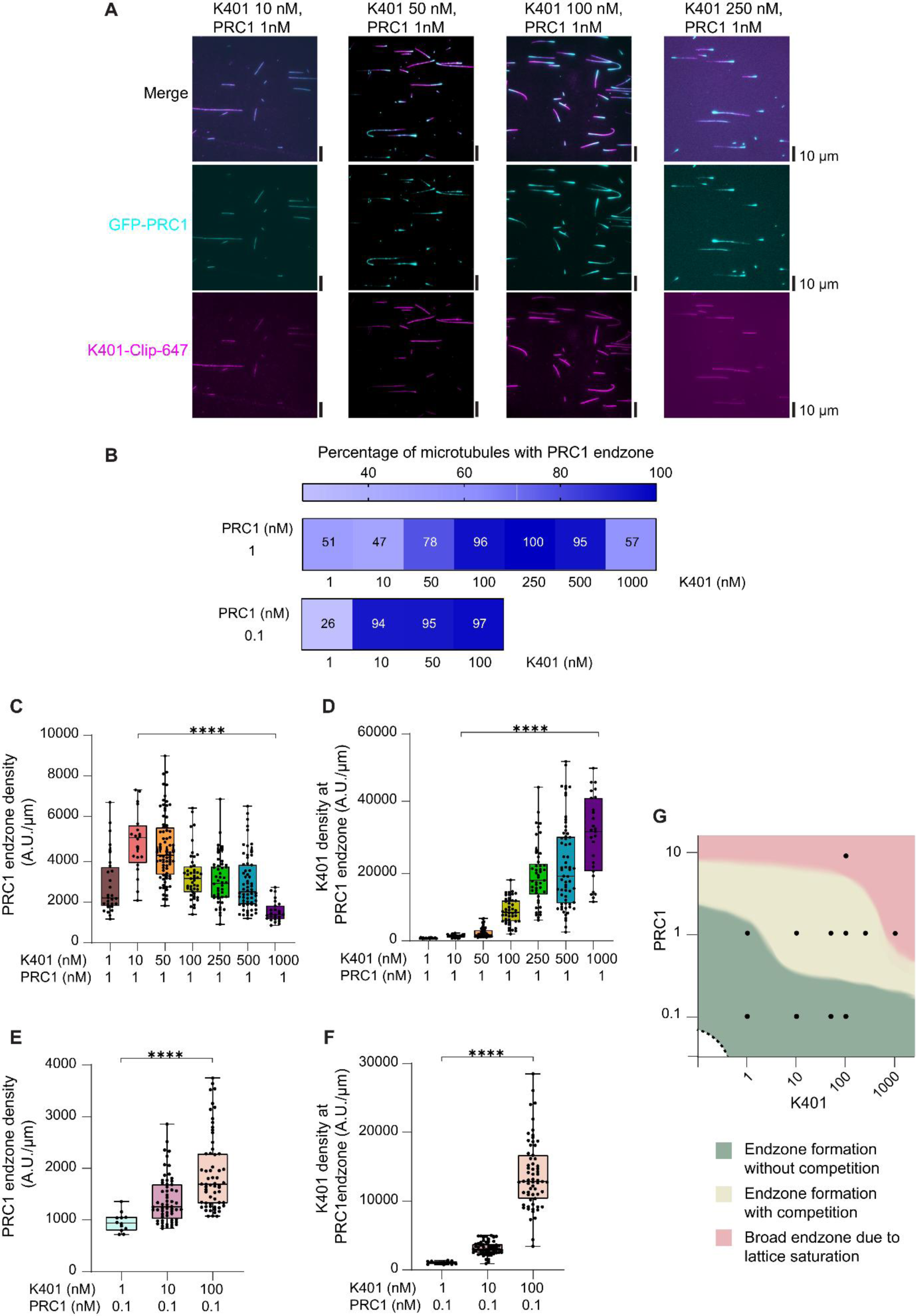
The relative concentration of K401 and PRC1 determines shepherding efficiency. (A) Representative fields of view (merged, GFP-PRC1 in cyan, K401-Clip-647 in magenta) from shepherding experiments at 1 nM GFP-PRC1 and 10-250 nM K401-Clip-647 at steady state. Scale bars, 10 μm. (B) The percentage of microtubules with PRC1 endzones at steady state for 1 nM GFP-PRC, and 1, 10, 50, 100, 250, 500, and 1000 nM K401-Clip-647 (N = 55, 43, 97, 48, 45, 66, 47). 0.1 nM GFP-PRC1 and 1, 10, 50, and 100 nM K401-Clip-647 (N = 38, 64, 86, and 65). Data re from three independent experiments, except the data for 1 nM PRC1 and 1000 nM K401 are from two independent experiments. Darker blue indicates greater shepherding efficiency. (C-D) Steady state endzone density of GFP-PRC1 (C) and K401-Clip-647 density at the PRC1 endzone (D) for 1nM PRC1 and 1, 10, 50, 100, 250, 500, and 1000 nM of K401-Clip-647. For PRC1 endzone density, the mean and standard deviation are 2821 ± 1440 (N=28), 4826 ± 1439 (N=20), 4569 ± 1567 (N=76), 3279 ± 1134 (N=46), 3009 ± 1152 (N=44), 2915 ± 1311 (N=62), 1533 ± 517 (N=26) A.U./µm, respectively. For K401 density, the mean and standard deviation are 836 ± 117 (N=28), 1471 ± 463 (N=20), 2494 ± 1369 (N=66), 8891 ± 3809 (N=46), 19284 ± 8577 (N=45), 21910 ± 12303 (N=63), and 30006 ± 11366 (N=27) A.U./µm, respectively. Data are from three independent experiments except the data for 1 nM PRC1 and 1000 nM K401 are from two independent experiments. Box edges, middle line, and whiskers are 25th and 75th percentiles, median and minimum and maximum values, respectively. P-values were calculated by unpaired nonparametric Kolmogorov-Smirnov t-test. (E-F) Steady state endzone density of GFP-PRC1 (E) and K401-Clip-647 density at the PRC1 endzone (F) for 0.1 nM PRC1 and 0.1, 10, 100 nM of K401. For PRC1 endzone density, the mean and standard deviation and N are 952 ± 187 (N=12), 1394 ± 460 (N=60), and 1911 ± 736 (N=61) A.U./µm, respectively. For K401 density, the mean and standard deviation are 1056 ± 210 (N=12), 3164 ± 966 (N=59), and 13862 ± 4971 (N=63) A.U./µm, respectively. The data are from three independent experiments. Box edges, middle line, and whiskers are 25th and 75th percentiles, median and minimum and maximum values, respectively. P-values were calculated by unpaired nonparametric Kolmogorov-Smirnov t-test. (H) Schematic summarizing the observed shepherding across experimental conditions. Shepherding creates endzones with increasing PRC1 density at 0.1 nM PRC1 and increasing amount of K401 motors (green region). However, upon increasing PRC1 to 1 nM, competition between PRC1 and K401 is observed in addition to shepherding, thereby reducing PRC1 endzone densities (yellow region). Less shepherding observed at high PRC1 or K401 concentrations potentially due to lattice saturation (red region). The plot depicts the approximate boundaries for protein shepherding based on experimental observations. The white area under the dashed curve represents conditions of low protein concentration that were not considered. Black dots represent experimentally tested concentrations.

In addition to the number of microtubules with endzones, the PRC1 intensity profile varied with K401 concentration. For example, at higher K401 concentration, the edge of the endzone appeared sharper but the fluorescence intensity was lower, visible in fluorescence images displayed with the same brightness and contrast (Fig. S9A). To quantify this, we measured the GFP-PRC1 and K401-Clip-647 density (intensity per unit length) in the endzone. Only filaments with endzones as determined by our criteria were analyzed (Supplementary Methods). For 1 nM PRC1, the PRC1 endzone density first increased with an increasing concentration of K401 from 1 to 10 nM and then decreased at higher K401 concentration (Fig. 6C). By contrast, the density of K401 motors in the PRC1 endzone increased with concentration as expected (Fig. 6D). These data suggest that binding competition between K401 and PRC1 may reduce the steady state PRC1 endzone density at higher K401 to PRC1 ratios. For the lower PRC1 concentration of 0.1 nM, the PRC1 endzone density increases as K401 concentration was increased from 1 to 100 nM (Fig. 6E). This suggests that reducing the concentration of PRC1 reduces the binding competition, allowing PRC1 density to increase over a larger K401 concentration range. As expected, the K401 endzone fluorescence density increased with K401 concentration (Fig. 6F). Consistent with the binding competition model, at fixed K401 concentration of 100 nM, the PRC1 endzone density increased with increasing PRC1 concentration, while the K401 endzone density decreased (Fig. S9C-E).

Thus, the interplay between the concentration of PRC1 and K401 tunes the shepherding efficiency along the microtubule and protein composition at endzones (Fig. 6G).

### Shepherding occurs for another MAP, the tetrameric CAPGly construct

Our model predicts that shepherding is possible for diffusive MAPs, independent of the molecular details of the microtubule-MAP interaction. To determine whether shepherding can occur for other motor-MAP combinations, we tested endzone formation of the cytoskeleton-associated protein-glycine-rich (CAPGly) domain which is found in many MAPs. The CAPGly domain differs from the microtubule-binding domain of PRC1, which is composed of a spectrin domain and an unstructured domain^16 40^. CAPGly domains interact with the C-terminal tail of α-tubulin which extends from the microtubule lattice^41^. We utilized an N-terminal construct of CLIP-170 (aa 1-481, previously referred to as the H2 construct^42^) which is dimerized via the coiled-coil and thus contains 4 CAPGly domains (hereafter 4xCAPGly-GFP). Previous work demonstrated that this 4xCAPGly-GFP construct binds diffusively along the microtubule lattice and does not accumulate at microtubule ends^42^. We paired this MAP with the rat kinesin-1 construct KIF5C(1-560) (rK560, Fig. S4F). Non-fluorescent rK560 and 4xCAPGly-GFP were incubated with immobilized GTP taxol-stabilized microtubules and ATP and imaged at steady state (Fig. 7A-C). 4xCAPGly-GFP accumulated at the ends of microtubules in endzones similar to those observed for PRC1. The endzones became narrower and denser with increasing rK560 concentration from 20 to 200 nM at a fixed concentration of 4xCAPGly-GFP. No endzones formed in the absence of ATP or rK560 (Fig. 7D,E), confirming that active motors were required for shepherding.

**Fig. 7.**
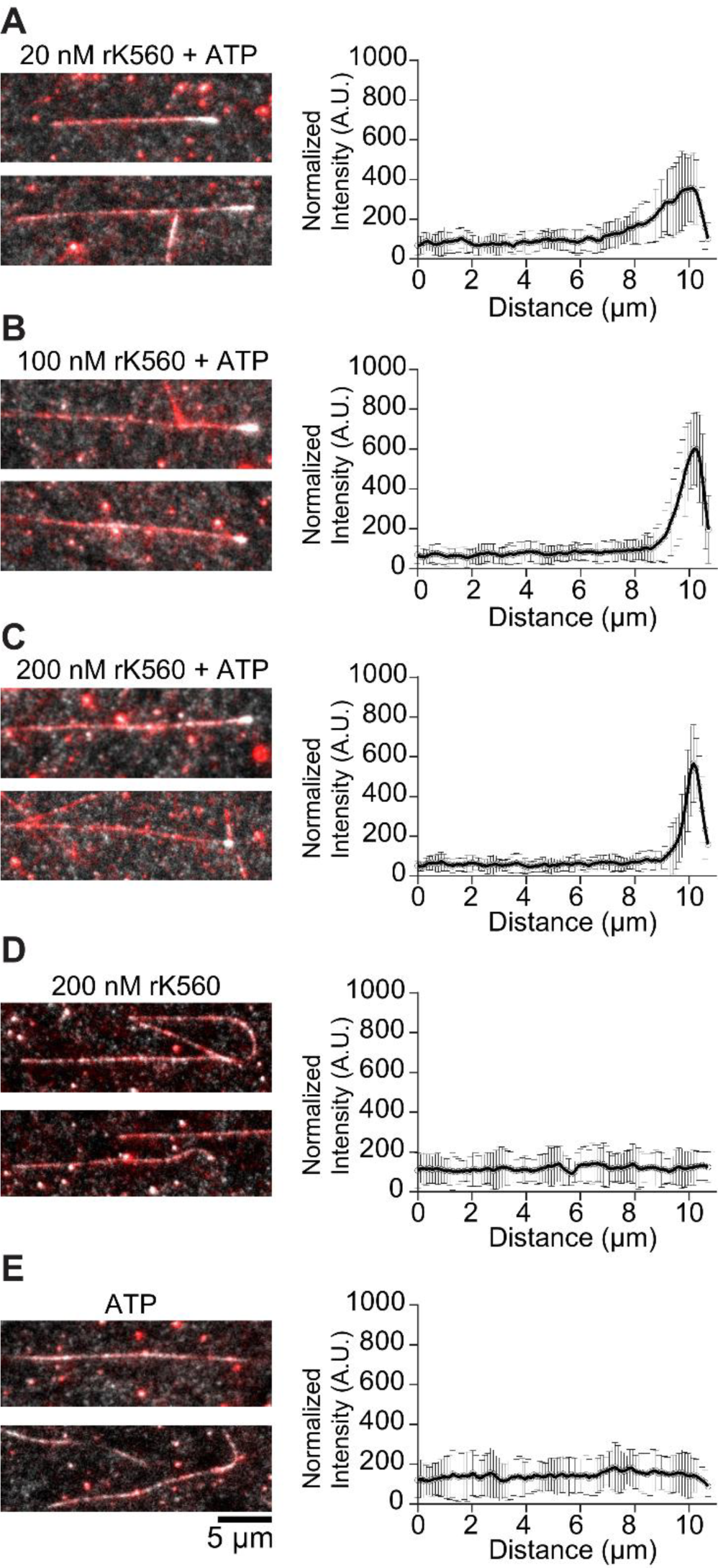
Shepherding with tetrameric CAPGly protein in the presence of KIF5C(1-560). Representative images (left) and fluorescence intensity distribution (right) for 0.74 nM 4xCAPGly-GFP (gray) bound to taxol-stabilized microtubules (red) with 20 nM K560 and ATP (A), 100 nM K560 and ATP (B), 200 nM K560 and ATP (C), 200 nM K560 (D), and ATP only (E). Fluorescence intensity presented as mean ± standard deviation, for 21-34 microtubules from two or three independent experiments.

## DISCUSSION

Our work identifies shepherding as a mechanism for self-organization of diffusive MAPs through rectified diffusion by multiple motors on a multi-protofilament lattice. Repeated steric interactions bias MAP diffusion, leading to directional transport and end accumulation of PRC1. The strength of the biased movement depends on geometry, MAP diffusion, and motor number: fewer accessible protofilaments, lower lateral diffusion, and higher motor number all enhance shepherding. Shepherding is thus a collective phenomenon that arises from the activity of multiple motors.

Shepherding provides a concrete experimental realization of rectified Brownian motion in an active matter system. This is conceptually related to Brownian ratchet models, where thermal fluctuations are rectified by a directional process to generate motion or force^43^. The MAP remains purely diffusive and does not consume chemical energy, yet directional motion emerges from interactions with active particles along the lattice. Motors act as an active bath, similar to models in which self-propelled particles rectify the motion of Brownian tracers or create directed tracer transport in ratchet-like landscapes^1,2,4^. Shepherding is thus part of a broader class of active lattice-gas phenomena for which steric exclusion and biased hopping generate non-trivial currents and spatial structure^44^. In shepherding, the driven and diffusive particles occupy a multi-lane microtubule lattice with varying protofilament accessibility. As a result, the microtubule can be tuned between quasi-one- and two-dimensional regimes, which controls how effectively steric encounters convert thermal motion into drift and end accumulation.

While high motor processivity is not required, shepherding crucially depends on diffusion of the MAP (Figs. 1,2). Shepherding occurred most effectively in our model for an intermediate “Goldilocks” value of the MAP longitudinal diffusion coefficient (Fig. 1). In the limiting case of an immobile MAP, no diffusion occurs. Even for a diffusive MAP, its diffusion may be restricted by crowding, limiting the potential for shepherding. For example, dense regions of PRC1 that crosslink microtubules in the spindle midzone would have limited capacity to be shepherded. In the other limit, if MAP diffusion is too high and it can switch protofilaments, then its movement is little affected by motors. Differences between diffusion of different types of MAPs can cause shepherding to sort them into different regions of the endzone (Fig. S2E).

In addition to diffusion, the microtubule-binding lifetime of the MAP is also an important determinant of shepherding. The average MAP displacement due to shepherding can be estimated as *d*_transport_ ≈ *v*_shepherd_*t*_bind_, where *v*_shepherd_ is the shepherding velocity and *t*_bind_ the MAP binding lifetime, which can be estimated analytically (Supplementary Material). For motor and MAP parameters based on K401 and PRC1 (Table S1) that lead to a MAP transport velocity of ∼100 nm/s, MAP lifetime must be ∼10 seconds to allow micron-scale MAP displacement. While shepherding transport velocity likely varies for different motor-MAP combinations, our model suggests that ≳ 1000 nm/*v*_shepherd_ is a rule of thumb for the minimum binding lifetime to easily observe shepherding transport. In addition, cooperative interactions that slow diffusion and increase MAP lifetime in clusters can help maintain the endzone accumulations that arise due to shepherding.

Conventional cargo transport by motors uses direct binding in which the cargo is tethered to the motor. Therefore, the motor and cargo move together at the same velocity and with similar processivity. In our results, MAP and motor velocities and lifetimes differ by approximately an order of magnitude or more, showing that MAP motion is not tied to individual motor trajectories. In contrast to conventional modes of transport, shepherding neither requires direct motor-cargo binding nor is it dependent on high motor processivity. This means that kinesins that are not typically considered as transport motors could drive shepherding. Because a number of non-motor MAPs act at the plus or minus ends of microtubules to regulate polymer dynamics, to crosslink microtubules, to form lateral bundles or asters, or to serve as platforms for recruiting other proteins^9–12^, the mechanism by which non-motor MAPs are transported to microtubule ends is of fundamental interest. A major class of mechanisms invoked for MAP transport is through specific motor-MAP interactions coupled with unidirectional long-distance transport by kinesins^20–25^. Shepherding is an alternative mode of directional movement and displacement of MAPs

Our results highlight the contribution of microtubule geometry and spatial variation of MAP diffusion to shepherding. While most previous modeling of protein motion on microtubules considers just a single protofilament, we found that for shepherding it was essential to model multiple protofilaments for quantitative agreement with experiments (Fig. 4). For shepherding by kinesin-1 motors that track along a single protofilament, MAP hopping between protofilaments lowers the transport velocity and average displacement because MAPs can hop out of the way of the motors (Fig. 1, Supplementary Material). Because of the importance of lateral diffusion, the number of accessible protofilaments on the microtubule plays a significant role in shepherding, with the highest velocity for fewer (nonzero) accessible protofilaments (Fig. 1). Even among accessible protofilaments, differences in lateral diffusion would be predicted to lead to differences in shepherding. For example, lateral hops across the seam of a microtubule would likely be slower than between other protofilaments, which would lead to faster shepherding of MAPs bound near the seam. Further, while in our experiments and model the MAPs were shepherded to microtubule plus ends, any location in a microtubule network with slower or obstructed diffusion could serve as an accumulation site for shepherded MAPs. For example, the transition of a crosslinking MAP from a single microtubule to a region of overlapping microtubules may stall shepherding due to reduced diffusion upon microtubule crosslinking. Similarly, lattice defects may act as a barrier to shepherding, and this may in turn promote repair or disassembly of microtubules from these sites. Therefore, shepherding could drive different modes of transport and sites of accumulation depending on the structure of both individual microtubules and the organization of the microtubule network, potentially offering a way to specify behavior according to the local environment. Shepherding could therefore help redistribute diffusive MAPs along long cellular microtubules, control MAP composition at plus ends, or maintain spatial asymmetry in dense or confined microtubule arrays.

In the future, it would be of interest to further understand what features of motors and MAPs allow shepherding. We have optimized the experiments to show this phenomenon most clearly, but such optimization and micron-scale transport are not required for shepherding to be relevant in cells, because even smaller-scale displacement can be important for MAP reorganization. Whether shepherding occurs in cells may be challenging to demonstrate conclusively, as definitive evidence requires ruling out binding of the transported MAP to any motor in the cell. We speculate that shepherding could be a mechanism to non-specifically “sweep” MAPs along microtubules in cells, limiting crowding and opening binding sites. We also note that shepherding and conventional transport are not mutually exclusive: if a motor and MAP bind for transport, shepherding could act to enhance the effective affinity of transport by continuing to transport cargo after unbinding events. Shepherding is also a possible mechanism to explain slow axonal transport, in which cargo movement occurs along axons at speeds low compared to motor speeds^45–48^. Selective shepherding of specific cargoes could be achieved through differences in their inherent diffusive properties or regulatory mechanisms like post-translational modifications. Notably, shepherding also does not require motor recycling from microtubule ends after cargo delivery, which would offer advantages in long axons. In addition, imaging of fluorescently tagged kinetochores in *Saccharomyces cerevisiae* lysates suggested that micron-scale movement of individual kinetochore molecules on microtubules may result from transient interactions with a kinesin motor, suggesting that it may be relevant in the context of mitosis^35^.

Other findings suggest that cargo diffusion and motor processivity can have unexpected contributions to transport. Recent work found that motors of only moderate processivity can effectively transport large cargos because the large, slowly diffusing cargo stays near the microtubule even after the motors unbind, increasing the effective processivity of transport ^49^. Similarly, the effective processivity of a kinesin motor can be enhanced by confinement in a narrow nuclear-envelope tether in fission yeast ^50^. These recent results therefore broaden our understanding of the molecular and physical mechanisms that enhance cargo transport in cells. Shepherding may extend beyond microtubules to other crowded polymer systems in cells where active and passive components coexist, such as in the spatial patterning of non-motor DNA-binding proteins for genome organization in the nucleus.

## ACKNOWLEDGEMENTS

This work was funded by NSF grant DMR1725065 and NIH grants 5R01GM124371 (MB) and 1R01GM155215 (RS), and the European Union’s Horizon Europe research and innovation programme under the Marie Skłodowska-Curie Actions, MSCA 101151485 (SAF). This work utilized the Summit supercomputer, which is supported by the National Science Foundation (awards ACI-1532235 and ACI-1532236), the University of Colorado Boulder, and Colorado State University. The Summit supercomputer is a joint effort of the University of Colorado Boulder and Colorado State University.

