## Supplemental methods and figures for "Shepherding by biased diffusion organizes diffusible proteins on microtubules"

### Supplemental material for
### Shepherding by biased diffusion organizes diffusible proteins on microtubules

Leila Farhadi, Shane A. Fiorenza, et al.

#### Computational model

Our model represents passive MAPs and active motors on microtubules as a lattice gas model. We utilize a tau-leaping kinetic Monte Carlo algorithm to evolve the system over time and reach steady-state. MAPs and motors are implicitly modeled in solution and are randomly drawn from their respective infinite reservoirs upon binding to microtubules [1].

##### Microtubules

Microtubules with multiple protofilaments are modeled as a two-dimensional lattice, where each ∼8-nm tubulin dimer corresponds to a discrete binding site. Each protofilament has $N = L/\delta$ binding sites, where $L$ is the length of the microtubule and $\delta$ is the size of a tubulin dimer. The total number of binding sites for a microtubule is therefore $N_{\mathrm{tot}} = N \times N_{\mathrm{pf}}$, where $N_{\mathrm{pf}}$ is the number of protofilaments in the microtubule. We assume that all protofilaments are aligned with one another and there are no offsets from the helicity of the microtubule. All binding sites have the same base binding affinity before binding interactions are taken into account. Microtubules are polar and have sites which correspond to a plus and minus end. The microtubule is assumed to be fixed in space and hydrodynamic interactions between bound proteins are neglected [1].

##### MAPs

Each MAP consists of one passive binding head that binds to the microtubule with rate $k_{\mathrm{on,p}} c_{\mathrm{p}} \tilde{N}_{\mathrm{tot}}$, where $k_{\mathrm{on,p}}$ is the per-site binding rate of MAPs, $c_{\mathrm{p}}$ is the bulk concentration of MAPs in solution, and $\tilde{N}_{\mathrm{tot}}$ is the number of unoccupied binding sites available. Bound MAPs unbind from the microtubule at rate $k_{\mathrm{off,p}}$ and, while bound, can diffuse longitudinally along the same protofilament or laterally to an adjacent protofilament with diffusion constants of $D_{\mathrm{long}}$ and $D_{\mathrm{lat}}$. Lattice hopping rates are obtained from diffusion constants via $k_{\mathrm{hop}} = 2D/\delta^2$. We assume that MAPs have short-range binding cooperativity that is represented as a nearest-neighbor interaction (range of one lattice site). The attractive potential has an energy of magnitude $\epsilon$ (in units of $k_{\mathrm{B}}T$). The cooperativity affects unbinding, giving

$$k_{\mathrm{on,p}} = k^0_{\mathrm{on,p}}, \tag{1}$$

$$k_{\mathrm{off,p}} = k^0_{\mathrm{off,p}} \exp[-n\epsilon], \tag{2}$$

where $n = 0, 1, 2, 3, 4$ is the number of nearest neighbors that are MAPs. We assume that MAPs and motors do not interact with one another through this potential. Parameter values and descriptions appear in Table S1; for additional details, see Ref. [1].

#### Motors

Each motor consists of two active binding heads connected by a rigid linkage that undergo an ATP hydrolysis cycle coupled to stepping. Motor heads can be bound to either ADP, ATP, ADP·Pi, or no ligand (empty). While in solution, both motor heads are assumed to be bound to ADP. The first motor head binds to the microtubule with rate $k_{\mathrm{on,m}} c_{\mathrm{m}} \tilde{N}_{\mathrm{tot}}$, where $k_{\mathrm{on,m}}$ is the per-site binding rate of motors, $c_{\mathrm{m}}$ is the bulk concentration of motors in solution, and $\tilde{N}_{\mathrm{tot}}$ is the number of unoccupied binding sites available. The first motor head to bind is always leading, i.e., closer to the plus-end than the unbound head. Upon binding to the microtubule, the first motor head releases ADP and becomes empty. While empty or ATP-bound, motor heads are assumed to be strongly bound and cannot unbind from the microtubule. Singly bound motors can therefore only unbind while in the "vulnerable" ADP·Pi-bound state. ATP binding to the empty head occurs at rate $k_{\mathrm{ATP}}$ and induces a conformational change that swings the second (unbound) head forward. The first (bound) head then hydrolyzes ATP to ADP·Pi at rate $k_{\mathrm{hydro}}$. Once ATP hydrolysis occurs, the motor can either unbind its first (bound) head at rate $k_{\mathrm{off,1}}$, terminating its run, or bind its second (unbound) head to the microtubule at rate $k_{\mathrm{on,m}} c_{\mathrm{eff}}$, making it doubly bound and continuing its run. Here $c_{\mathrm{eff}}$ is the effective concentration of the unbound motor head when constrained to be near the microtubule by the bound head. The ratio between $k_{\mathrm{on,m}} c_{\mathrm{eff}}$ and $k_{\mathrm{off,1}}$ determines how many steps occur on average before the motor fully unbinds and thus sets the average single-molecule motor processivity

$$\delta n_{\mathrm{cycles}} = \delta \frac{k_{\mathrm{on,m}} c_{\mathrm{eff}}}{k_{\mathrm{off,1}}}. \tag{3}$$

The singly bound unbinding rate $k_{\mathrm{off,1}}$ is used to set motor processivity in our model. Importantly, motor processivity can be significantly altered by lattice crowding, as motors will be stuck in the vulnerable singly bound state if the site immediately in front of them is occupied and the second head cannot bind. While the motor is doubly bound, the front head cannot unbind and the rear head unbinds with rate $k_{\mathrm{off,2}}$. Upon the rear head unbinding, the motor is again in the singly bound state, having advanced forward one site along the microtubule. The time to complete one full ATP hydrolysis cycle sets the average single-molecule velocity of motors, given by:

$$\frac{\delta}{t_{\mathrm{cycle}}} = \delta \left[ \frac{1}{k_{\mathrm{ATP}}} + \frac{1}{k_{\mathrm{hydro}}} + \frac{1}{k_{\mathrm{on,m}} c_{\mathrm{eff}}} + \frac{1}{k_{\mathrm{off,2}}} \right]^{-1}. \tag{4}$$

Hydrolysis of ATP is the rate-limiting step of the cycle, and thus we use $k_{\mathrm{hydro}}$ to set motor velocity. Parameter values and descriptions appear in Table S1; for additional details, see Ref. [1].

#### Steric exclusion

No two binding heads can occupy the same binding site at the same time. Any steric effects that do not occur on the microtubule lattice are neglected, e.g., the trailing head of a singly bound motor can always swing forward during a conformational change regardless of crowding.

#### End effects

Motor end pausing is implemented by disabling the conformational change induced by ATP binding when a motor head is bound to the microtubule plus end. The ATP hydrolysis cycle continues as usual other than this, and the trailing head can still re-bind to the microtubule as long as the site behind the motor remains unoccupied. We also assume that MAPs cannot diffuse off either end of the microtubule.

| | Description | Value | Notes |
|---|---|---|---|
| **General** | | | |
| $k_\mathrm{B}T$ | Thermal energy | 4.1 pN·nm | Room temperature |
| $t$ | Total simulation time | $100 - 600$ s | Typical experiment time |
| $t_\mathrm{s}$ | Data snapshot interval | 0.1 s | Time between data output |
| $dt$ | Timestep | $5 \times 10^{-6}$ s | Time increment of each kMC step |
| **Microtubules** | | | |
| $L$ | Length | $8 - 40$ $\mu$m | Experimental values |
| $\delta$ | Site size | 8.2 nm | Ref. [2] |
| $N_\mathrm{pf}$ | Protofilament number | 1-8 | Reference value of 8, varied to explore model; see text |
| **MAPs** | | | |
| $\epsilon$ | Interaction energy | 0.6 $k_\mathrm{B}T$ | Fit to match experiment |
| $n_\mathrm{max}$ | Maximum neighbors | 4 | Geometry of 2D lattice |
| $k^0_\mathrm{on,p}$ | Per-site binding rate | 0.0024 nM$^{-1}$s$^{-1}$ | Increased by a factor of 10 from Ref. [3] |
| $c_\mathrm{p}$ | Bulk concentration | 0.1 or 1.0 nM | Experimental values |
| $k^0_\mathrm{off,p}$ | Off rate | 0.01 s$^{-1}$ | Estimated from experiment |
| $D_\mathrm{long}$ | Longitudinal diffusivity | 0.13 $\mu$m$^2$s$^{-1}$ | Ref. [4] |
| $D_\mathrm{lat}$ | Lateral diffusivity | 0.66 $\mu$m$^2$s$^{-1}$ | Fit to match experiment |
| **Motors** | | | |
| $k_\mathrm{on,m}$ | Per-site binding rate | 0.0036 nM$^{-1}$s$^{-1}$ | Increased by a factor of 10 from Ref. [3] |
| $c_\mathrm{m}$ | Bulk concentration | $0 - 100$ nM | Experimental values |
| $c_\mathrm{eff}$ | Effective concentration of second unbound head | $1 \times 10^6$ nM | Adjusted from Ref. [3] to keep binding-to-unbinding ratio while singly bound constant |
| $k_\mathrm{ATP}$ | ATP binding rate | 5000 s$^{-1}$ | 2 mM ATP, Ref. [5] |
| $k_\mathrm{hydro}$ | ATP hydrolysis rate | 75 s$^{-1}$ | Sets single-molecule velocity to 600 nm/s, consistent with experimental measurements |
| $k_\mathrm{off,1}$ | Singly bound off rate | 20 s$^{-1}$ | Sets single-molecule processivity to 1.2 $\mu$m, consistent with experimental measurements |
| $k_\mathrm{off,2}$ | Doubly bound off rate | 2400 s$^{-1}$ | Ref. [5] |

Table S1: Parameter values used in simulation. Additional details in Refs. [1, 3]

#### Simulation algorithm

Each simulation timestep is implemented using a hybrid tau-leaping algorithm [6], which samples from the binomial and Poisson distributions to predict the number of kinetic Monte Carlo (kMC) events that will occur in a given timestep. The binomial distribution is sampled for events with a constant probability throughout the simulation, such as ATP hydrolysis by a motor head or protein binding (these probabilities are constant in time and identical for every site in the absence of any long-range coupling) [7]. For events with a probability that varies, we sample the Poisson distribution. This can occur, for example, when protein unbinding is force dependent, or if cooperative interactions give different binding kinetics at different lattice sites. In this case, we compute the pairwise partition function of each object that an event can act on to calculate the average number of events expected to occur in the timestep [7]. We then sample the Poisson distribution to choose the number of events that occur. Events are executed in random order on randomly selected members of the appropriate population. Multiple events can target the same object, e.g., binding of the second head and unbinding of the first head for a singly bound motor. We enforce that no two events can act on the same object in the same timestep, a good approximation if the timestep is sufficiently small. Each kMC substep implements the following steps:

1. Loop over all active objects and sort into appropriate populations.

2. If appropriate, check binding equilibration status of each protein species. This is done by finding the average number of proteins bound for each species in a time window $t_c$. If the change in number bound between two windows is less than twice their standard deviations added in quadrature, the protein species is considered equilibrated. Once all species are equilibrated, the simulation is considered equilibrated.

3. Sample the appropriate statistical distribution for each possible event. If the event has a constant probability, use the binomial distribution. If the event has a probability that can change in space or time, sum all relevant pairwise partition functions to calculate the expected average and sample the Poisson distribution.

4. If two or more events target the same object, discard at random until only one remains. The probability of each event determines the relative weight when sampling to discard.

5. Execute each event in a random order on random members from the appropriate population.

#### MAP and motor motility measurements

Velocity measurements in simulation exclude data from runs that end less than one micron away from the microtubule end. This is done to ensure long-lived protieins in the dense endzone do not bias the average. For simulation data that appear in Fig. 4, this exclusion region is extended to three microns away from the microtubule plus end since the concentrations used lead to increased endzone accumulation. Displacement and lifetime measurements use all data, regardless of where the run ended. Displacement is defined as the difference between the final and initial bound positions of the proteins, and lifetime is defined as the time from binding to unbinding. Averages of these quantities are determined by the arithmetic mean of many events in the simulations. Flux is obtained by averaging the number of proteins that cross the midpoint of the microtubule toward the plus and minus end at each timestep. Reported net flux values are calculated by subtracting the average minus-end-directed flux from the average plus-end-directed flux.

#### Simulated fluorescence images

To generate simulated kymographs, the MAP fractional occupancy is averaged for 1 second to mimic image collection time. We then apply a Gaussian filter to the average MAP occupancy to create the green channel of the image. The same method is used with motor occupancy data to create the red and blue channels, making motors appear as purple. A padding region of 15 pixels is added to both sides of the microtubules. This creates a one-dimensional row of intensity data, which is used as a single horizontal slice in kymographs. The final image is created by arranging many of these horizontal slices in a two-dimensional matrix. Within each individual phase diagram, all kymographs are generated using the same simulated fluorescence intensity per molecule. The simulated fluorescence intensity per molecule is varied between phase diagrams with different protein concentration to make molecules visible.

##### Reproduction of previous results

To check our model implementation, we ran simulations using the same ingredients and parameters as previous work [8]. Our model utilizes an ATP hydrolysis cycle to facilitate motor stepping, and average motor off rate depends on local crowding as discussed above. If the site in front of the motor is occupied, the second head cannot unbind and the motor spends more time in the vulnerable singly bound state, reducing overall lifetime in crowded environments [1]. Single-molecule data in this and other studies that this is the case for K401. Because the site in front of motors is often occupied by a MAP during shepherding, simulations of our model using the same effective concentration and single-molecule off rate as previous work do not achieve a similar steady state (Fig. S1A). In our model, this effect can be partially offset by reducing both the singly bound off rate and effective binding concentration of the second head by the same factor, maintaining overall single molecule processivity but reducing sensitivity to crowding (Fig. S1B,C). For a more direct comparison with previous work, we implemented the same motor stepping model in CyLaKS, in which motors consist of a single binding head that has a fixed probability to advance forward one site while bound. With this implementation, our simulations reproduce the results of previous work [8] as expected (Fig. S1D).

#### Analytic model

Building on related work in driven lattice gases [9, 10, 11, 12, 8, 13, 14, 15, 16, 17], we developed an analytic model of shepherding. On a multi-protofilament lattice, the microtubule has $N_{\text{pf}}$ parallel lanes (protofilaments) of $N$ longitudinal sites and lattice spacing $\delta = 8.2$ nm. Motors (species A for active) step longitudinally on a single protofilament with rate $k_+$ to the right and corresponding motor velocity $v_a = k_+\delta$. The bound motor site occupancy is $\rho_a$. MAPs (species P for passive) hop on the lattice with rates $k_\parallel = D_\parallel/\delta^2$ longitudinally (either direction) and $k_\perp = D_\perp/\delta^2$ laterally between protofilaments. The bound MAP density is $\rho_p$.

The parameter values estimated from simulation-experiment comparison and shown in table S1 are $D_\parallel \approx 0.13$ $\mu$m$^2$/s, $D_\perp \approx 5D_\parallel$, $v_a \approx 600$ nm/s corresponding to rates $k_\parallel \approx 1930$ s$^{-1}$, $k_\perp \approx 9670$ s$^{-1}$, $k_+ \approx 73$ s$^{-1}$. Note that the MAP hopping rate exceeds the motor stepping rate. Motors and MAPs interact only via steric exclusion, so that no site is occupied by more than one particle of either type. Both species also bind/unbind from solution. The lattice occupancy away from the endzone of motors for 10 nM bulk motor concentration is $\rho_a = (k^a_{\text{on}}c)/(k^a_{\text{on}}c + k^a_{\text{off}}) = (0.0036 \times 10)/(0.036 + 0.685) = 0.050$. For 0.1 nM PRC1 concentration we estimate $\rho_p = (0.0024 \times 0.1)/(0.00024 + 0.01) = 0.023$.

##### Single protofilament shepherding velocity

We want to predict the longitudinal drift velocity $v_{\text{shep}}$ of a MAP in the bulk of the lattice away from the endzone on a single protofilament. The interactions that cause biased MAP diffusion are steric exclusion with a motor either just behind or just ahead of the MAP. These are determined from the occupancy of a motor one site behind or one site ahead of the MAP

$$p_- = P(n_a^{i-1} = 1 \mid n_p^i = 1), \tag{5}$$
$$p_+ = P(n_a^{i+1} = 1 \mid n_p^i = 1). \tag{6}$$

Here the occupancy $n^i_{a,p}$ is 1 if the site is occupied and 0 if it is empty. In these configurations, the only MAP movement that can occur is to hop forward (when a motor is behind) or backward (if a motor is ahead). Therefore once we know $p_\pm$ the drift velocity is

$$v_{\text{shep}} = \delta k_\parallel (p_- - p_+) \tag{7}$$

To determine $p_-$, consider the relative position of a motor and closest MAP, separated by $m \geq 0$ empty sites. If the motor and MAP are adjacent then $m = 0$. The gap changes if the MAP hops forward (rate $k_\parallel$, increasing $m$), the MAP hops backward (rate $k_\parallel$, decreasing $m$), or the motor steps forward (rate $k_+$, decreasing $m$). At $m = 0$ only the forward MAP hop is available. If we assume that the region ahead of the MAP is dilute, then a forward hop is essentially always allowed and the gap undergoes random walk on $m \geq 0$ with forward rate (increasing $m$) $k_\parallel$ and backward rate $k_\parallel + k_+$, with reflecting boundary conditions at $m = 0$. We assume the probability distribution $P(m) = P_m$ of gap size reaches steady state, since the rates (especially of MAP hopping) are fast compared to the shepherding velocity.

The master equations for $P_m$ are

$$\frac{dP_0}{dt} = (k_\parallel + k_+)P_1 - k_\parallel P_0, \tag{8}$$

$$\frac{dP_m}{dt} = k_\parallel P_{m-1} - (2k_\parallel + k_+)P_m + (k_\parallel + k_+)P_{m+1} \qquad m > 0. \tag{9}$$

At steady state, this gives a recursion relation

$$P_m = r^m P_0, \qquad r = \frac{k_\parallel}{k_\parallel + k_+}. \tag{10}$$

After using normalization to set $P_0 = 1 - r$, we have

$$P_m = (1 - r)\, r^m. \tag{11}$$

The configuration with a motor one site behind the MAP occurs with probability

$$P_0 = \frac{k_+}{k_\parallel + k_+}, \tag{12}$$

with mean gap $\langle m \rangle = k_\parallel / k_+$. For our parameters ($k_\parallel = 1930$ s$^{-1}$, $k_+ = 73$ s$^{-1}$) this is $P_0 = 0.036$, with the motor otherwise within $\langle m \rangle \approx 26$ sites.

This calculation leading to equation (12) assumes a motor is present behind the MAP. When this occurs is set by the slower binding kinetics. We assume that after any motor binds behind the MAP, it catches up at rate $k_+$ and stays in the steady-state distribution until it unbinds at rate $k^a_{\text{off}}$. The next motor is typically $1/\rho_a$ sites back and catches up in time $\sim 1/(k_+\rho_a)$. The fraction of time the MAP has a nearby motor is therefore

$$f = \frac{1/k^a_{\text{off}}}{1/k^a_{\text{off}} + 1/(k_+\rho_a)} = \frac{k_+\rho_a}{k_+\rho_a + k^a_{\text{off}}}. \tag{13}$$

Because the gap random walk reaches steady state quickly compared with binding turnover, binding and the random walk can be treated as independent. As a result, the typical motor occupancy one site behind the MAP is

$$p_- = f\, P_0 = \frac{k_+\rho_a}{k_+\rho_a + k^a_{\text{off}}} \frac{k_+}{k_\parallel + k_+}. \tag{14}$$

In principle, we should also calculate $p_+$, the occupancy of the site ahead of the MAP, but typically this will be low. The site ahead is governed by the same random walk where a motor ahead of the MAP steps forward freely at $k_+$ while the MAP chasing it hops forward at $k_\parallel$, so the gap to a leading motor has forward rate $k_+ + k_\parallel$ and backward rate $k_\parallel$, biased away from contact. On average, a MAP does not catch a motor ahead of it, so the site ahead is occupied only by a new motor binding, $p_+ \approx k^a_{\rm on} c/k_+ \ll \rho_a$, and is negligible.

Neglecting $p_+$ the shepherding velocity is

$$v_{\rm shep} \approx \delta k_\parallel p_- = \frac{k_+\rho_a}{k_+\rho_a + k^a_{\rm off}} \frac{\delta\, k_\parallel k_+}{k_\parallel + k_+}. \tag{15}$$

The second factor is the pair velocity

$$v_{\rm pair} = \frac{\delta\, k_\parallel k_+}{k_\parallel + k_+}, \tag{16}$$

the rate at which a coupled motor-MAP pair advances one site by alternating a MAP hop and a following motor step. The shepherding velocity is then $v_{\rm shep} = f v_{\rm pair}$, which means the MAP moves at the pair velocity for the fraction of time $f$ that a motor is near it.

The shepherding velocity in equation (15) has two regimes set by the binding factor $f$. When motor occupancy is high ($k_+\rho_a \gg k^a_{\rm off}$), $f \to 1$ and $v_{\rm shep} \to v_{\rm pair}$ because the MAP almost always has a nearby motor. When motor occupancy is low ($k_+\rho_a \ll k^a_{\rm off}$), $f \to k_+\rho_a/k^a_{\rm off}$ and $v_{\rm shep} \to v_{\rm pair}\, k_+\rho_a/k^a_{\rm off}$, linear in motor flux. The pair velocity itself crosses over between the two single-particle rates. When MAP hopping is rate-limiting ($k_\parallel \ll k_+$), $v_{\rm pair} \to \delta k_\parallel = D_\parallel/\delta$, the MAP diffusive hopping rate. When motor stepping is rate-limiting ($k_+ \ll k_\parallel$), $v_{\rm pair} \to \delta k_+ = v_a$, the motor velocity. In our experiments $k_+ \approx 73$ s$^{-1}$ is small compared to $k_\parallel \approx 1930$ s$^{-1}$, so motor stepping is the slower step and $v_{\rm pair}$ lies closer to $v_a$ than to $D_\parallel/\delta$.

For our parameters ($\delta = 8.2$ nm, $k_\parallel = 1930$ s$^{-1}$, $k_+ = 73$ s$^{-1}$, $\rho_a = 0.050$ for 10 nM motors, and $k^a_{\rm off} = 0.685$ s$^{-1}$ from the 1.46 s motor lifetime),

$$P_0 = k_+/(k_\parallel + k_+) = 0.036, \tag{17}$$
$$f = k_+\rho_a/(k_+\rho_a + k^a_{\rm off}) = 3.65/4.33 = 0.84, \tag{18}$$
$$v_{\rm pair} = \delta k_\parallel k_+/(k_\parallel + k_+) = 8.2 \text{ nm} \times 1930 \times 73/2003 \text{ s}^{-1} = 577 \text{ nm/s}, \tag{19}$$
$$v^{\rm 1PF}_{\rm shep} = f\, v_{\rm pair} = 0.84 \times 577 \text{ nm/s} = 485 \text{ nm/s}. \tag{20}$$

Our single-protofilament simulations for this concentration found $v_{\rm shep} \sim 400$ nm/s, a reasonable $\sim 20\%$ discrepancy with this simplified model.

#### Multi-protofilament extension

To extend to a multi-protofilament lattice, we focus on the limit of fast lateral hopping $k_\perp \gg k_+$. In this limit the MAP redistributes laterally across all $N_{\rm pf}$ accessible protofilaments on a timescale short compared with motor stepping, so motors are effectively static while the MAP samples protofilaments and the MAP-motor encounter dynamics can be averaged over protofilaments. This causes two main changes to the calculation. The effective time for a motor to move close to a MAP decreases, and the fraction of time that a MAP spends near a motor on a given protofilament decreases.

On a multi-protofilament lattice, motors step on each of $N_{\rm pf}$ protofilaments in parallel. Because the MAP can hop laterally to the protofilament carrying the incoming motor on a timescale fast compared with motor stepping, this increases the effective approach rate by $N_{\rm pf}$. The binding factor is then

$$f_N = \frac{N_{\rm pf} k_+\rho_a}{N_{\rm pf} k_+\rho_a + k^a_{\rm off}}, \tag{21}$$

which reduces to the single-protofilament result at $N_{\rm pf} = 1$ and saturates near 1 when $N_{\rm pf} k_+ \rho_a \gg k^a_{\rm off}$.

The pair velocity $v_{\rm pair} = \delta k_\parallel k_+/(k_\parallel + k_+)$ describes a MAP-motor pair advancing one site by alternating a MAP hop and a motor step. This occurs when the MAP is on the same protofilament as the motor. With fast lateral hopping the MAP samples accessible protofilaments uniformly, so the fraction of the time the MAP is on the motor protofilament is $1/N_{\rm pf}$.

Combining the enhanced binding factor and the reduction in protofilament residency gives a multi-protofilament shepherding velocity

$$v_{\rm shep} = \frac{f_N}{N_{\rm pf}} v_{\rm pair} = \frac{k_+ \rho_a}{N_{\rm pf} k_+ \rho_a + k^a_{\rm off}} v_{\rm pair}. \tag{22}$$

For $N_{\rm pf} = 1$, $f_N = f$ and $v_{\rm shep} \to f v_{\rm pair}$, recovering the single-PF result. For $N_{\rm pf} k_+ \rho_a \gg k^a_{\rm off}$, $f_N \to 1$ and $v_{\rm shep} \to v_{\rm pair}/N_{\rm pf}$.

For our parameters with $N_{\rm pf} = 8$, $\rho_a = 0.050$, $k_+ = 73$ s$^{-1}$, $k^a_{\rm off} = 0.685$ s$^{-1}$, and $v_{\rm pair} = 577$ nm/s, $f_8 \approx 0.977$, and $v_{\rm shep} \approx 70.5$ nm/s. The formula predicts a monotonic decrease of $v_{\rm shep}$ with $N_{\rm pf}$ in our parameter regime which qualitatively matches the trends in the multi-protofilament simulations (main text fig. 1G). This analytic calculation slightly overestimates the shepherding velocity by 20-50% compared to simulation results.

The derivation assumes fast lateral hopping such that motors are effectively static on the timescale of the MAP lateral diffusion across accessible protofilaments. This requires $\delta^2/D_\perp \ll 1/k_+$, or $D_\perp \gg k_+ \delta^2 = v_a \delta \approx 5 \times 10^{-3}\,\mu$m$^2$/s. Our $D_\perp = 0.65\,\mu$m$^2$/s exceeds this by two orders of magnitude. As $D_\perp$ falls, the MAP becomes effectively confined to one protofilament on the relevant timescale and the single-protofilament formula takes over.

##### Endzone steady state profile

Once we have $v_{\rm shep}$, we can compute the mean-field steady-state MAP density profile in the endzone from a drift-diffusion equation with reflective plus-end boundary. In the bulk, the MAP current is

$$J = v_{\rm shep} \rho_p - D_\parallel \partial_x \rho_p. \tag{23}$$

In steady state with $J = 0$, a reflective plus end with no flux through the right boundary,

$$\rho_p(x) = \rho_p^\infty \exp\left[\frac{v_{\rm shep}}{D_\parallel}(x - L)\right] = \rho_p^\infty\, e^{(x-L)/\ell}, \quad \ell = \frac{D_\parallel}{v_{\rm shep}}. \tag{24}$$

Here $L$ is the plus-end position and $\rho_p^\infty$ is the bulk MAP density far from the end. This exponential profile is the mean-field solution; correlations at high density are important and determine the actual endzone profile which deviates significantly from the exponential form. However, the decay length $\ell$ provides an analytic estimate of endzone size. It scales inversely with shepherding velocity. For $D_\parallel = 0.13\,\mu$m$^2$/s and $v_{\rm shep} \approx 80$ nm/s, $\ell = 1.6\,\mu$m. This is comparable to the endzone width $\sim$ 1-3 $\mu$m measured experimentally.

#### Supplementary Methods

##### Cloning

K401-BIO-His6 construct contains the first 401 residues of *Drosophila* kinesin heavy chain fused to 87-residue region of *Escherichia coli* biotin carboxyl carrier protein (BCCP) followed by a six-histidine tag to facilitate purification [18]. The K401-Clip-His6 was generated by replacing the

BCCP with a CLIP tag sequence.

For the PRC1 constructs, Human-PRC1-isoform-1 cDNA was inserted into the bacterial expression vector pET-DUET containing a Tobacco Etch Virus (TEV) protease cleavable N-terminal His-tag as described before [19, 20]. To construct the N-terminal-GFP tagged PRC1, eGFP sequence was inserted between the TEV cleavage site and the N-terminus of PRC1 with a three amino acid linker(‘AAA’). To generate a Snap-PRC1 construct, SNAP sequence was inserted between the TEV cleavage site and the N terminus of PRC1.

The rKIF5C(1-560)-halo-2xstreptagII construct was prepared by fusing the truncated KIF5C(1-560) motor with halo tag and 2xstreptagII tags by restriction digestion and subcloning into the pFastBac1 vector.

The 6xHis-4xCAPGly-mEGFP constructs were prepared by PCR amplification of DNA corresponding to amino acids 3 – 484 of Rat CLIP1 (Uniprot Q9JK25), followed by a C-Terminal mEGFP tag. PCR amplification was performed with Prime STAR Max DNA Polymerase (Takara bio. Cat# R045B) with the following forward primers for 4xCAPGly: 5’ – GCGGCAGCCATATGCTCGA GCTGAAACCCAGCGGGCTGAAG – 3’. The same reverse primer was used to amplify sequence: 5’ – CTTTCGGGCTTTGTTAGCAGCCGTTACTTGTACAGCTCGTCC ATGC – 3’. The C-terminally mEGFP-tagged CLIP1 fragments were joined with a BAMHI-linearized pET15b vector via Gibson assembly (NEB).

#### Protein purification

K401: The K401-Clip and K401-BIO-H6 proteins were expressed in Rosetta™(DE3)pLysS (Novagen) *Escherichia coli*. The constructs were expressed overnight at 20°C after induction with 1 mM IPTG. Cells were lysed by brief sonication in a buffer containing 50 mM phosphate (pH 8), 300 mM NaCl, 1 mM MgCl2, 30 mM imidazole, 5% glycerol, 0.15 % Tween, 40 $\mu$M ATP, 2 mM TCEP, 1% Igepal, 0.2 mg/mL lysozyme, HALT protease inhibitor cocktail, 1.4 mM PMSF, 2 mM Benz-HCL, and Benzonase. The lysate was clarified by centrifugation and the supernatant was incubated with Ni-NTA resin (QIAGEN) for 1 hr. The mixture was poured into a column, washed with 50 mM phosphate (pH 8), 300 mM NaCl, 1 mM MgCl2, 30 mM imidazole, 5% glycerol, 0.15% Tween, 40 $\mu$M ATP, 0.5 mM TCEP. Protein was eluted with a buffer containing 50 mM phosphate (pH 7), 200 mM NaCl, 1 mM MgCl2, 400 mM imidazole, 5% glycerol, 40 $\mu$M ATP, and 0.5 mM TCEP. Peak fractions were pooled and purified further by size exclusion chromatography (Superose 6 -10-300 GL) in a gel filtration buffer with 50 mM phosphate (pH 7), 300 mM NaCl, 1 mM MgCl2, 5% glycerol, 40 $\mu$M ATP, and 10 mM $\beta$-mercaptoethanol. After concentration, sucrose was added to make the final 30% sucrose solution, then it was frozen in liquid nitrogen, and stored at -80°C.

PRC1: All the PRC1 constructs were expressed in Rosetta™(DE3)pLysS (Novagen) *Escherichia coli* and purified using procedures similar to those described previously before [19, 20]. The constructs were expressed for 1 hour, after induction with 1 mM IPTG. Cells were lysed by brief sonication in a buffer containing 50 mM phosphate (pH 8), 300 mM NaCl, 30 mM imidazole, 5% glycerol, 0.15% Tween, 2 mM TCEP, 1% Igepal, 0.2 mg/mL lysozyme, and HALT protease inhibitor cocktail, 1.4 mM PMSF, 2 mM Benz-HCL, and Benzonase. The lysate was clarified by centrifugation and the supernatant was incubated with Ni-NTA resin (QIAGEN) for 1 h. The mixture was poured into a column, washed with 50 mM phosphate (pH 8), 300 mM NaCl, 30 mM imidazole, 5% glycerol, 0.15% Tween, 0.5 mM TCEP. Then it eluted with a buffer containing 50 mM phosphate (pH 7), 150 mM NaCl, 400 mM imidazole, 5% glycerol, and 0.5 mM TCEP. Peak fractions were pooled and purified further by size exclusion chromatography (Superose 6 -10-300 GL) in a gel filtration buffer with 50 mM phosphate (pH 7), 500 mM NaCl, 5% glycerol, and 10 mM $\beta$-mercaptoethanol. After concentration, sucrose was added to make the final 30% sucrose solution, then it was frozen in liquid nitrogen, and stored at -80°C.

For Snap and Clip protein labeling, purified proteins were incubated with SNAP-Surface® 647 or CLIP-Surface™647 or CLIP-Surface™488 in a 1:5 (protein:dye) molar ratio, at room temperature for 5 -10 minutes, followed by incubation at 4°C overnight. The unbound dye was removed by repeated dilution and centrifugation through an Amicon Ultra-15 Centrifugal Filter Unit (Millipore Sigma), prior to size exclusion chromatography. A comparison of the absorbance of labeled protein at 650 nm or 488 nm with 280 nm yielded a labeling efficiency of 39%, 34%, and 21.8% per monomer for Snap-647-PRC1, K401-Clip-647, and K401-Clip-488 protein preparations, respectively.

CAPGly-mEGFP: 6x-His tagged CAPGly-mEGFP constructs were transformed and expressed in BL21-CodonPlus -RILC competent cells (Aligent Technologies Cat# 230245). Cells were grown in Terrific Broth to an OD600 of 0.6 at 37°C, then induced with 0.5 M IPTG for 18 hours at 18°C. Cells were pelleted and stored at -80°C until purification. 4 x CAPGly construct were purified similar to the H2 construct in Bieling et al., 2008. For purification, cells were resuspended in lysis buffer (50 mM KPi (pH 7.5), 500 mM NaCl, 1 mM MgCl2, 1 mM $\beta$-mercaptoethanol, 1 mM PMSF, 1.0 mg/ml lysozyme (Sigma-Aldrich, Cat# L6876), Benzonase nuclease (Sigma-Aldrich, Cat# E1014), and 1 x protease inhibitor cocktail (SIGMAFAST, Sigma-Aldrich, Cat# S8830). Bacterial lysates were sonicated and clarified via centrifugation. Clarified lysates were applied to Ni-NTA resin that was pre-equilibrated with lysis buffer. After binding of the protein, the column was washed with 10 column volumes of low imidazole wash buffer (50 mM KPi (pH 7.5), 500 mM NaCl, 1 mM MgCl2, 8.5 mM Imidazole (pH 7.4), and 1 mM $\beta$-mercaptoethanol) and 3 column volumes of high imidazole wash buffer (50 mM KPi (pH 7.5), 500 mM NaCl, 1 mM MgCl2, 125 mM Imidazole (pH 7.4), and 1 mM $\beta$-mercaptoethanolE). Proteins were eluted from the Ni-NTA resin with elution buffer (50 mM KPi (pH 7.5), 500 mM NaCl, 1 mM MgCl2, 300 mM Imidazole (pH 7.4), and 1 mM $\beta$ME). Each protein was then gel filtered over a HiLoad 16/600 Superdex 200 pg column (Cytiva) equilibrated with 50 mM KPi (pH 7.5), 150 mM NaCl, 1 mM MgCl2, and 1 mM $\beta$ME. Pooled fractions containing each protein were concentrated with appropriately sized MWCO centrifugal concentrators (MilliporeSigma), aliquoted, flash frozen, and stored at -80°C until use. Protein concentrations were determined via Bradford protein assay.

rKIF5C(1-560)-halo-2xstreptagII: Sf9 cells obtained from Thermo Fisher Scientific were cultured in suspension with serum-free sf900 II SFM medium (Thermo Fisher Scientific) supplemented with antibiotic antimycotic (Gibco) in flasks at 28°C in a non-CO2 nonhumidified incubator with an orbital shaker platform set at 110 rpm. The cells were infected with baculovirus generated according to the Bac-to-Bac system (Invitrogen). In brief, plasmids encoding truncated-tagged motors in the pFastBac1 vector were transformed into DH10Bac Escherichia coli to generate recombinant bacmids. Bacmid DNA was isolated with the HiPure Plasmid DNA miniprep kit (Invitrogen) and confirmed by PCR analysis. Recombinant bacmid DNA was transfected into Sf9 cells using Cellfectin II (Invitrogen) to produce the P1 recombinant baculovirus. 7d after transfection, the supernatant containing P1 baculovirus was collected by centrifugation at 3,000 rpm for 3 minutes at 4°C. The baculovirus was amplified by successive infection of Sf9 cells to generate P2 and P3 baculoviruses. Baculovirus-containing supernatants were stored at 4°C in the dark. To purify protein, Sf9 cells were infected with 3% P3 baculovirus (vol/vol). 3 d after infection, the cells were harvested by centrifugation for 15 minutes at 3,000 rpm at 4°C. The pellet was washed once with PBS and resuspended in ice-cold lysis buffer (200 mM NaCl, 4 mM MgCl2, 0.5 mM EDTA, 1 mM EGTA, 0.5% igepal, 7% sucrose, and 20 mM imidazole-HCl, pH 7.5) supplemented with 2 mM

ATP, 1 mM PMSF, 5 mM DTT, and protease inhibitor cocktail. After 30 minutes incubation on ice, the lysates were clarified by ultracentrifugation for 20 minutes at 20,000 rpm in F12-8x50y rotor (Sorvall 3421), and the supernatants were incubated with strep-Tactin beads (Strep-Tactin XT 4Flow resin, Iba) for 1h at 4°C with rotation. The supernatant from beads bound proteins were drained in a PD-10 column and washed with wash buffer (150 mM KCl, 25 mM imidazole-HCl, pH 7.5, 5 mM MgCl2, 1 mM EDTA, and 1 mM EGTA) supplemented with 1 mM PMSF, 3 mM DTT, 3 mM ATP, and protease inhibitor cocktail. Bound proteins were eluted by elution buffer (25 mM KCl, 25 mM imidazole-HCl, pH 7.5, 5 mM EGTA, 2 mM MgCl2, 2 mM DTT, 0.1 mM ATP, 1 mM PMSF, protease inhibitor cocktail and 10% glycerol) supplemented with 50 mM biotin in 6x0.5 mL fractions. SDS PAGE of the eluted fractions was done and pure fractions were selected by looking at the bands. Pure protein fractions were combined and dialyzed in dialysis buffer (25 mM imidazole-HCl, pH 7.5, 25 mM KCl, 5 mM EGTA, 2 mM MgCl2, 2 mM DTT, 0.1 mM ATP and 10% glycerol). After 2 hours the buffer was changed to a fresh dialysis buffer and dialyzed overnight to remove biotin from the sample at 4°C. The protein sample was collected by centrifugation, and aliquots were snap frozen in liquid nitrogen and stored in -80°C until further use.

KIF4A: Full length human KIF4A containing a TEV protease cleavable C-terminal His-tag was expressed from a pFastBac1 plasmid in sf9 cells and purified as previously described [20].

#### Microtubule polymerization

For experiments with GFP-PRC1 and/or K401-Clip-647: GMPCCP polymerized and taxol-stabilized biotinylated X-rhodamine labeled microtubules were prepared as described previously [21, 3]. Briefly, GMPCPP seeds were prepared by mixing unlabeled bovine tubulin, X-rhodamine-tubulin, and biotin tubulin (Cytoskeleton), in BRB80 buffer (80 mM PIPES pH 6.8, 1.5 mM MgCl2, 0.5 mM EGTA, pH 6.8) and GMPCPP. The mixture was spun at 90000 rpm for 5 minutes at 4°C to clarify. Then, the supernatant was aliquoted and flash frozen and stored at -80°C. To polymerize microtubules, the tube that contains GMPCPP seed was transferred to a 37°C heating block and was incubated for 3-4 hours, while protected from light exposure. Then, 100 $\mu$l of warm BRB80 buffer was added to the microtubules and spun at 75000 rpm for 10 minutes at 37°C to remove free unpolymerized tubulin. Following the centrifugation step, the supernatant was discarded, and the pellet was washed by a round of centrifugation with 100 $\mu$l BRB80 supplemented with 20 $\mu$M taxol. The pellet was resuspended in BRB80 containing 20 $\mu$M taxol and stored at 28°C covered in foil.

For experiments that study the dynamics of PRC1 or K401 at single molecule resolution (Fig. 4, 5A-E, Fig. S6): GMPCCP polymerized, and taxol-stabilized biotinylated HiLyte Fluor™ 488 labeled microtubules were prepared similar to the protocol discussed above.

For an experiment that studies the dynamics of PRC1 and K401 at single molecule resolution (Fig. 5H): GMPCCP polymerized, and taxol-stabilized biotinylated X-rhodamine labeled microtubules were prepared similar to the protocol discussed above.

For microtubule gliding experiments: GMPCCP polymerized, and taxol-stabilized X-rhodamine labeled microtubules were prepared. GMPCCP seeds were prepared by mixing unlabeled bovine tubulin, X-rhodamine-tubulin (Cytoskeleton) diluted in BRB80 buffer, and GMPCCP. The microtubules were polymerized as described above.

For experiments with rKIF5C and/or CAPGly: HiLyte647-labeled microtubules were polymerized from purified tubulin including 6% Hily647-labeled tubulin (Cytoskeleton) in BRB80 buffer (80 mM Pipes/KOH pH 6.8, 1 mM MgCl2, and 1 mM EGTA) supplemented with 1 mM GTP and

2.5 mM MgCl2 at 37°C for 30 minutes. 20 $\mu$M taxol in prewarmed BRB80 buffer was added and incubated at 37°C for additional 30 minutes to stabilize microtubules. Microtubules were stored in the dark at room temperature for further use.

#### Pull-down assay

The pull-down assays between GFP-PRC1 and K401-Clip or KIF4A (Fig. 3) were performed using GFP-beads (ChromoTek; gta-10) and the following buffer: BRB80, 0.5 mM MgCl2, 0.5 mM EGTA, 5% sucrose, 50 mM KCl, 0.1% Triton X-100, 100 $\mu$M ATP, 1x Halt protease inhibitors, 1 mM TCEP and 1 mM DTT.

To reduce non-specific binding of K401-Clip and KIF4A to GFP-beads, we included a pre-clearing step. Each motor was first incubated with 10 $\mu$l of ChromoTek bab-20 beads. These beads are equivalent to GFP-beads of ChromoTek gta-10, but they lack the crosslinked nanobody that recognizes the GFP tag. This step eliminates any nonspecific binding to GFP-beads. After incubation, samples were centrifuged. Background proteins were trapped to ChromoTek bab-20 beads, while supernatants with motors were collected and used in subsequent steps for the pull-down assay.

GFP-beads (ChromoTek; gta-10) were equilibrated with pull-down buffer, followed by incubation with GFP-PRC1 (150 nM) for 1 hour at 4°C. Next, the beads were washed two times with pull-down buffer to remove unbound GFP-PRC1, followed by incubation with 2.5 or 5 $\mu$M K401-Clip or 0.5 $\mu$M KIF4A for 1 hour at 4°C. The beads were washed five times with pull-down buffer, resuspended with SDS-PAGE gel-loading buffer, and boiled. Eluted complexes were loaded onto 4-20 Novex 4-20% Tris-Glycine gel (Life Technologies; XP04200BOX) and stained with Coomassie Brilliant Blue. SDS-PAGE gels were scanned using the Odyssey scanning system (LICOR). The pull-down experiment was performed two times with 5 $\mu$M K401-Clip and once with 2.5 $\mu$M K401-Clip. Control experiments with 0.5 $\mu$M KIF4A were conducted three times. All experiments showed the same result. Non-specific binding of K401-Clip and KIF4A to GFP-beads was tested using the same protocol in the absence of GFP-PRC1.

#### TIRFM assays

Flow chamber preparation for experiments with PRC1 and/or K401 (Fig. 3, 4, 5, 6, S4, S5, S6, S8, S9): The microscope slides (Gold Seal Cover Glass, 24 × 60 mm, thickness No. 1.5) and coverslips (Gold Seal Cover Glass, 18 × 18 mm, thickness No. 1.5) were cleaned and functionalized with biotinylated PEG and non-biotinylated PEG, respectively, to prevent nonspecific surface sticking, as previously described [22]. A paper cutter (Silhouette, Portrait 3) was used to cut three flow chambers on double-sided adhesive sheets (Soles2dance, 9474-08x12 - 3M9474LE 300LSE). Flow chambers were assembled using a coated slide and coverslip separated by the pre-cut double-sided adhesive sheets. Each flow chamber volume was approximately 10 $\mu$l.

Endzone formation assays with GFP-PRC1 and K401-Clip-647: Fluorescently labeled biotinylated microtubules were immobilized in a flow chamber using 0.2 mg/ml neutravidin. After 10 minutes of incubation, the free filaments were washed out using 2X volume of assay buffer (BRB80 buffer supplemented with 0.5 mM MgCl2, 0.5 mM EGTA, 5% sucrose, and 5 mM TCEP) with 20 $\mu$M Taxol. GFP-PRC1 and K401-Clip-647 proteins were diluted to the required concentration in the assay buffer supplemented with 1 mg/ml BSA and 1 $\mu$M ATP. Lastly, GFP-PRC1 and K401-Clip-647 were flowed into the imaging chamber in assay buffer supplemented with 1 mM ATP, 10 mM Trolox, 0.5 $\mu$M k-casein, and 20 $\mu$M Taxol. For all experiments, oxygen scavenging mix comprising of 40 mg/ml glucose oxidase, 20 mg/ml catalase, 800 mM glucose, and 0.5% $\beta$-mercaptoethanol was included in the final buffer. Timelapse sequence of images was acquired either immediately or when the system reached steady state as indicated.

Dynamics of PRC1 molecules at single molecule resolution (Fig. 4, Fig. S6): Endzone formation assays were performed using a mixture of Snap-647-PRC1 and non-fluorescent PRC1 with non-fluorescent K401 motors. The ratios of Snap-647-PRC1 and unlabeled PRC1 were determined empirically as 10% (Fig. 4A-B, Fig. S6A, C-D, Movie 2, Movie 3), 34% (Fig. 4C), 1.4% (Fig. 4D-E), and 0.5% (Fig. S6E) of fluorescently labeled PRC1 molecules. For these experiments, we used GMPCCP polymerized, and taxol-stabilized biotinylated HiLyte Fluor™ 488 labeled microtubules to minimize crosstalk between microtubule and protein channels.

Dynamics of K401 molecules at single molecule resolution (Fig. 5A-E, Fig. S8): Endzone formation assays were performed using a mixture of K401-Clip-647 and non-fluorescent K401-Clip with non-fluorescent PRC1 molecules. Mixing K401-Clip-647 and non-fluorescent K401-Clip molecules resulted in 0.3% (Fig. 5A-B), 0.08% (Fig. 5C-D), and 0.1% (Fig. 5E) of fluorescent K401-Clip-647 motors. We prepared GMPCCP polymerized, and taxol-stabilized biotinylated HiLyte Fluor™ 488 labeled microtubules to minimize crosstalk between microtubule and protein channels.

Dynamics of K401 and PRC1 molecules at single molecule resolution (Fig. 5H): Endzone formation assays were performed using a mixture of K401-Clip-488, non-fluorescent K401-Clip, Snap-647-PRC1, and non-fluorescent PRC1 molecules. Mixing fluorescently labeled with non-fluorescent proteins resulted in 0.03% and 1.9% of fluorescent K401-Clip-488 and fluorescent Snap-647-PRC1 proteins.

Microtubule gliding assay with GFP-PRC1 and K401 (Movie 1): Biotinylated K401 molecules ( 180 nM) were immobilized on the glass coverslip using a 0.2 mg/ml neutravidin. Then, the flow chamber was washed with assay buffer (BRB80 buffer supplemented with 0.5 mM MgCl2, 0.5 mM EGTA, 5% sucrose, and 5 mM TCEP). GMPCCP polymerized, and taxol-stabilized biotinylated X-rhodamine labeled microtubule solution was flushed into the flow chamber and incubated for a few minutes. Next, 1 nM GFP-PRC1 and 190 nM of a mixture of K401 and K401-Clip-647 was flowed into the flow chamber in assay buffer supplemented with 1 mg/ml BSA, 1 $\mu$M ATP, 0.5 $\mu$M k-casein, 20 $\mu$M Taxol, 10 mM Trolox, and oxygen scavenging mix. Time lapse images were subsequently acquired to visualize microtubule gliding at steady state.

Endzone formation experiments with 4xCAPGly-GFP and/or rKIF5C (Fig. 7): A flow chamber ( 10 $\mu$l volume) was assembled by attaching a clean #1.5 coverslip (Fisher Scientific) to a glass slide (Fisher Scientific) with two strips of melted parafilm. Polymerized microtubules were diluted in BRB80 buffer supplemented with 10 $\mu$M taxol and infused into flow chamber. The microtubules were then incubated for 3 minutes to allow for nonspecific adsorption to the coverslips. Subsequently, blocking buffer (1 mg/ml casein in P12 buffer (12 mM Pipes/KOH pH 6.8, 2 mM MgCl2, and 1 mM EGTA)) was infused and incubated for another 3 minutes. Finally, 740 pM 4xCAPGly protein, along with or without the indicated concentration of rKIF5C(1-560) protein and 2 mM ATP in the motility mixture (0.3 mg/ml casein, 10 $\mu$M taxol, and oxygen scavenging solution (1 mM DTT, 1 mM MgCl2, 10 mM glucose, 0.2 mg/ml glucose oxidase, and 0.08 mg/ml catalase in P12 buffer) were added to the flow cells. The flow chamber was then sealed with molten paraffin wax.

#### Microscopy

Experiments with GFP-PRC1 and/or K401-Clip-647 (Fig. 3, 6, Fig. S4, S5, S9): Experiments were performed on a total internal reflection fluorescence (TIRF) inverted microscope (Nikon Eclipse Ti microscope) equipped with a 100x oil/1.45 NA objective (Nikon, Plan Apo $\lambda$) and an Andor iXon 512x512 pixel EM camera. The system has a 488 nm laser (Coherent Sapphire 488 LP, 150 mW), a 561 nm laser (Coherent Sapphire 561 LP, 150 mW), and a 640 nm laser (Coherent CUBE 640-100C, 100 mW). Filter cubes for each laser line was selected to minimize crosstalk. Image acquisition was controlled using $\mu$Manager (7) 2.0 and all assays were performed at room temperature [23, 24].

Experiments that study dynamics of PRC1 or K401 molecules at single molecule resolution (Fig. 4, 5A-E, S6, and S8): Experiments were performed on a total internal reflection fluorescence (TIRF) Nikon Ti-E inverted microscope equipped with a Ti-ND6-PFS perfect focus system and an APO TIRF 100x oil/1.49 NA objective (Nikon). The system has an EMCCD camera (Andor iXon Ultra, DU-897U-CSO-#BV), and a 488 nm, 561 nm and 640 nm laser. Filter cubes for each laser line were selected to minimize crosstalk (Chroma, TRF49904 (488 nm), TRF49909 (561 nm), and TRF49914 (640 nm)). Image acquisition was controlled using Nikon Elements software and all the assays were performed at room temperature.

Experiments that study dynamics of PRC1 and K401 molecules at single molecule resolution (Fig. 5H): Experiments were performed on a total internal reflection fluorescence (TIRF) Nikon Eclipse Ti2 inverted microscope equipped with a perfect focus system, an APO TIRF 100x oil/1.49 NA objective (Nikon), and iLas2 ring TIRF module. The system has an EMCCD camera (Andor iXon Ultra, DU-897U-CSO-#BV), and a 488 nm, 561 nm and 640 nm laser. Filter cubes for each laser line were selected to minimize crosstalk (Chroma, TRF49904 (488 nm), TRF49909 (561 nm), and TRF49914 (640 nm)). Image acquisition was controlled using Nikon Elements software and all the assays were performed at room temperature.

Experiments with rKIF5C and CAPGly (Fig. 7): Images were captured by TIRF microscopy using an inverted microscope Ti-E/B (Nikon) equipped with the perfect focus system (Nikon), a 100x oil/1.49 NA TIRF objective (Nikon), three 20-mW diode lasers (488 nm, 561 nm, and 640 nm) and an electron-multiplying charge-coupled device detector (iXon X3DU897; Andor Technology). Image acquisition was controlled using Nikon Elements software and all the assays were performed at room temperature.

#### Analysis

Software: Fiji (NIH) was used to process the image files. From the images, individual microtubules were selected and kymographs generated by MultipleKymograph plug-in (`http://fiji.sc/Multi_Kymograph`) (Fig. S5). These kymographs were used by a MATLAB code for further quantifications. To visualize the movements of individual PRC1 or K401 molecules, we used KymoResliceWide plug-in (`https://imagej.net/KymoResliceWide`) to generate kymographs (Fig. 4, 5, S6). All the plots and statistical analysis in the manuscript were generated using GraphPad Prism except otherwise stated. Schematic cartoons are made using BioRender and Adobe illustrator.

Time evolution of GFP-PRC1 and K401-Clip-647 distributions on microtubule (Fig. S4, S5): A custom MATLAB script was used to measure microtubule length and fluorescence intensity of GFP-PRC1 and K401-clip-647 along the microtubule at each time point from the kymographs (`https://github.com/lfarhadi/Shepherding-SubramanianLab`). Fluorescence density was calculated as fluorescence intensity per unit length at each time point. The GFP-PRC1 density on microtubule versus time plot was fitted into an exponential function to calculate a characteristic binding time for each filament and reported as mean $\pm$ standard deviation. Reported protein density vs time results are from three independent experiments. The PRC1 density at 1 nM GFP-PRC1 and K401 density at 100 nM K401-Clip-647 are calculated from two independent experiments.

GFP-PRC1 endzone analyses (Fig. 6, S9): To determine the percentage of microtubules with endzones at steady state, we performed fluorescence line scan analysis using Fiji to obtain GFP-PRC1 fluorescence intensity along each microtubule. We established the following criterion to score a microtubule as one having an endzone: (1) the fluorescence signal should have an absolute maximum value towards one end of the microtubule and (2) the width of the GFP-PRC1 peak at the microtubule end should be less than half of total microtubule length. This criterion counts only the filaments with one distinct peak in PRC1 fluorescence line scan at the microtubule end. Hence, the filaments with multiple peaks, peaks in the middle of filament, or gradient in their PRC1 signal without a clear peak were not counted as microtubules with PRC1 endzone. The percentage of filaments with an endzone from this analysis are presented in Fig. 6 and S9.

For microtubules that met the endzone criteria, PRC1 and K401 endzone density was extracted from the fluorescence intensity profiles of GFP-PRC1 and K401-Clip-647 obtained from the kymographs using a custom MATLAB script (`https://github.com/lfarhadi/Shepherding-SubramanianLab`). First, the GFP-PRC1 fluorescence intensity profile on microtubules was used to determine the endzone position and length (full width at half maximum, FWHM), and then endzone GFP-PRC1 density. Next, K401-Clip-647 intensity profile corresponding to the endzone position and length, determined from the GFP-PRC1 imaging, was used to determine K401-Clip-647 density (Fig. 6, S9). The results of these quantifications are displayed in Box-Whisker plots using GraphPad Prism. Box edges, middle line, and whiskers are 25th and 75th percentiles, median and minimum and maximum values, respectively. The mean and standard deviation are listed for GFP-PRC1 and K401-Clip-647 endzone density for different experimental conditions. Data from three independent experiments were analyzed, except the data for 1 nM GFP-PRC1 and 1000 nM K401-Clip-647 that is from two independent experiments.

To avoid fluorescence signal saturation, the experiments in Fig. 6, S9 with 0.1, 1, 10 nM GFP-PRC1, had to be performed at different laser powers (30%, 20%, and 10% respectively). We used background fluorescence intensities at these three different excitation laser powers to determine normalized fluorescence intensity at endzone. This allowed for comparison between GFP-PRC1 endzone densities under different assay conditions as shown in Fg. S6C-D.

CapGly endzone quantifications (Fig. 7): The fluorescence intensity of CAPGly protein along microtubules was analyzed using Fiji. A line (length = 100 pixels, width = 3 pixels) was drawn from the microtubule lattice to the microtubule end along filaments. The fluorescence intensity distribution of CAPGly protein along individual microtubules was normalized by subtracting the minimum fluorescence intensity value on the same microtubules. Plots were generated using GraphPad Prism. Fluorescence intensities are reported as mean $\pm$ standard deviation, for 21-34 microtubules from two or three independent experiments.

Data analysis of single molecule imaging experiments: PRC1 mean velocity (Fig. 4, S6): We generated kymographs for each microtubule from a five-minute-long imaging timeseries using KymoResliceWide plugin, FIJI. For each filament, we used a macro code in Fiji (`https://forum.image.sc/t/line-selection-tool-for-kymograph-velocity-measurement/32723`) to measure the slope of Snap-647-PRC1 trajectories to determine the mean transport velocity. Histograms and scatter plots of Snap-647-PRC1 mean velocities were generated and fitted to lognormal distributions using GraphPad Prism from four independent experiments per condition. The geometric mean and geometric standard deviation of Snap-647-PRC1 velocity is listed for each condition.

K401 mean velocity and processivity (Fig. 5, S8): We used KymographClear and KymographDirect2.1 software tools7 for K401-Clip-647 movement analysis. The KymographClear, a macro for ImageJ, generated kymographs from two-minute-long imaging timeseries. Then, KymographDirect, a stand-alone tool, extracted microtubule-bound lifetime and velocity from individual trajectories extracted from the kymographs. The Histograms of K401-Clip-647 lifetime were fitted into an exponential function, and the characteristic time with 95% confidence interval from the fit is reported. Histograms of K401-Clip-647 mean velocity were fitted to normal distributions and the mean ± standard deviation is reported. Data was analyzed from three independent experiments per condition.

Displacement and MSD analysis for PRC1 molecules (Fig. S7): This analysis was challenging and limited by the long lifetimes and diffusive movement of PRC1 molecules, which frequently cause PRC1 trajectories to cross each other or merge. This severely constrains the length of detected trajectories by the software, where the detected duration of the tracks is much shorter than the lifetime of PRC1 molecules. Bearing these limitations in mind, we used KymographClear and KymographDirect2.1 software tools, which were the best among different softwares tested, for automated movement analysis of Snap-647-PRC1. We analyzed 2 and 2.5-minute-long imaging timeseries from 0.1 nM PRC1 and 0.1 nM PRC1 and 100 nM K401, respectively. KymographClear generates kymographs from the timeseries data, and KymographDirect returns position vs time of PRC1 trajectories. For 0.1 nM PRC: the median length of detected trajectories is 6.8 ± 22.8 s from N=136 trajectories. For 0.1 nM PRC and 100 nM K401: the median length of detected trajectories is 11.3 ± 25.7 s from N=157 trajectories and two independent experiments.

Histograms of PRC1 displacement are displayed for 0.17, 0.83, 1.16, 1.66, 2.49, 3.32 s time intervals at 0.1 nM PRC1, and 0.1 nM PRC1 and 100 nM K401. The distributions were fitted to normal distribution, then mean displacement, and SEM were extracted for each condition (Fig. S7A-C). Mean squared displacement (MSD) vs time is also calculated and plotted in (Fig. S7D). Because the tracks for single PRC1 molecules generated by the software are short and not representative of the data, the fit parameters derived from the data are unreliable. However, a qualitative examination of the data shows significant differences in PRC1 displacement between the absence and presence of K401.

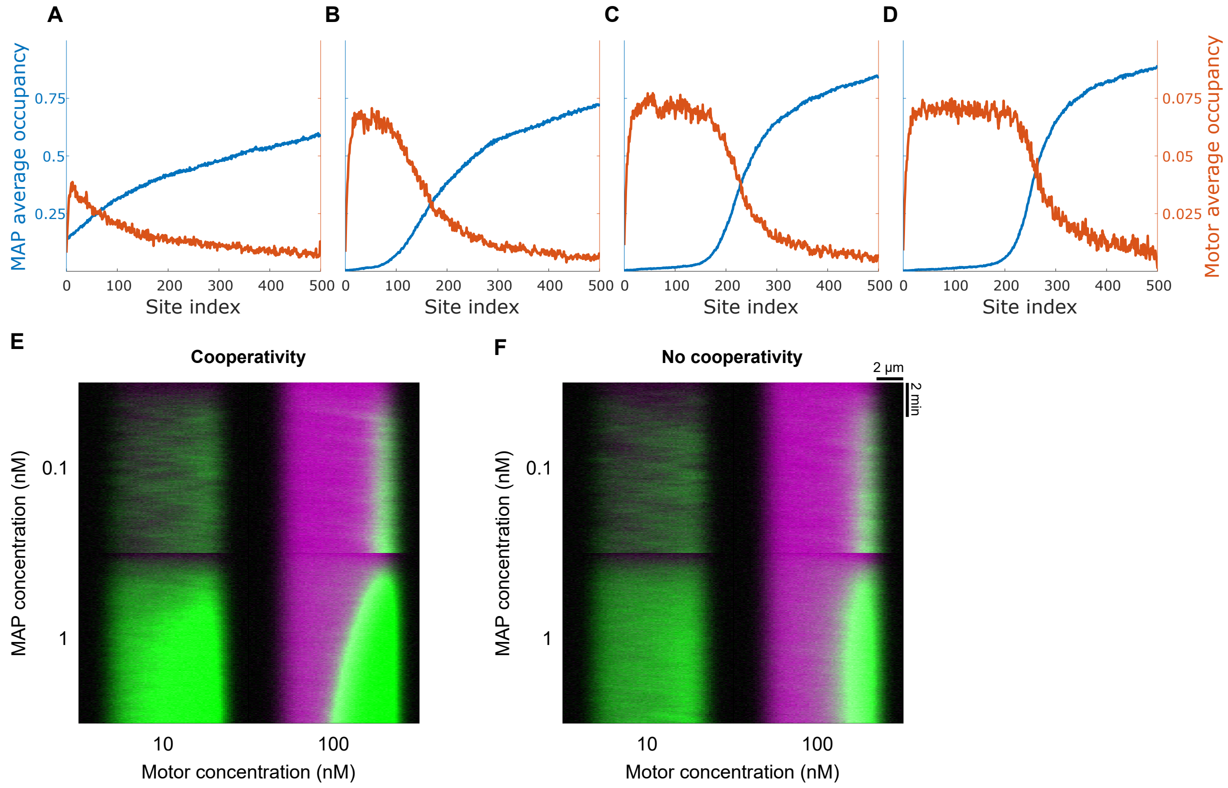


Fig. S1: **Reproduction of previous simulation results and effects of cooperativity on endzone accumulation.** (A-D) Comparison of our simulations to results of previous work [8]. In all panels, motor and MAP concentration were set to 33.52 nM and 0.504 nM, respectively to match per-site binding rate, and MAP unbinding rate and diffusivity were set to 0.0017 $s^{-1}$ and 0.0269 $\mu m^2 s^{-1}$ to match off-rate and hopping rate used in previous work. Average MAP occupancy (left axis, blue) and motor occupancy (right axis, orange) as a function of site along the filament for parameters with (A) motor singly bound unbinding rate 465 $s^{-1}$ and ATP hydrolysis rate 12.131 $s^{-1}$ to match single molecule motor lifetime and velocity of previous work. (B, C) Effective concentration $c_{\text{eff}}$ and singly-bound off-rate $k_{\text{off},1}$ reduced by a factor of (B) 10 and (C) 100 so that single molecule lifetime remains the same, but motors have increased lifetime when the site ahead of them is obstructed. ATP hydrolysis rate of (B) 12.513 $s^{-1}$ and (C) 18.255 $s^{-1}$ was used to maintain the single-molecule velocity. In (A-C), other parameters values are the values of Table S1. (D) Results of a motor model without an ATP hydrolysis cycle. As in previous work, motors are represented as a single binding head that has a fixed rate of hopping forward one site. Parameters were motor per-site on-rate, 0.12 $s^{-1}$, off-rate, 1.56 $s^{-1}$, and stepping rate 12 $s^{-1}$ as in previous work. (E, F) Simulated kymographs showing accumulation of MAPs at the microtubule endzone with (E) and without (F) short-range cooperativity between MAP molecules. Microtubules are 1000 sites $\approx$ 8 $\mu$m long.

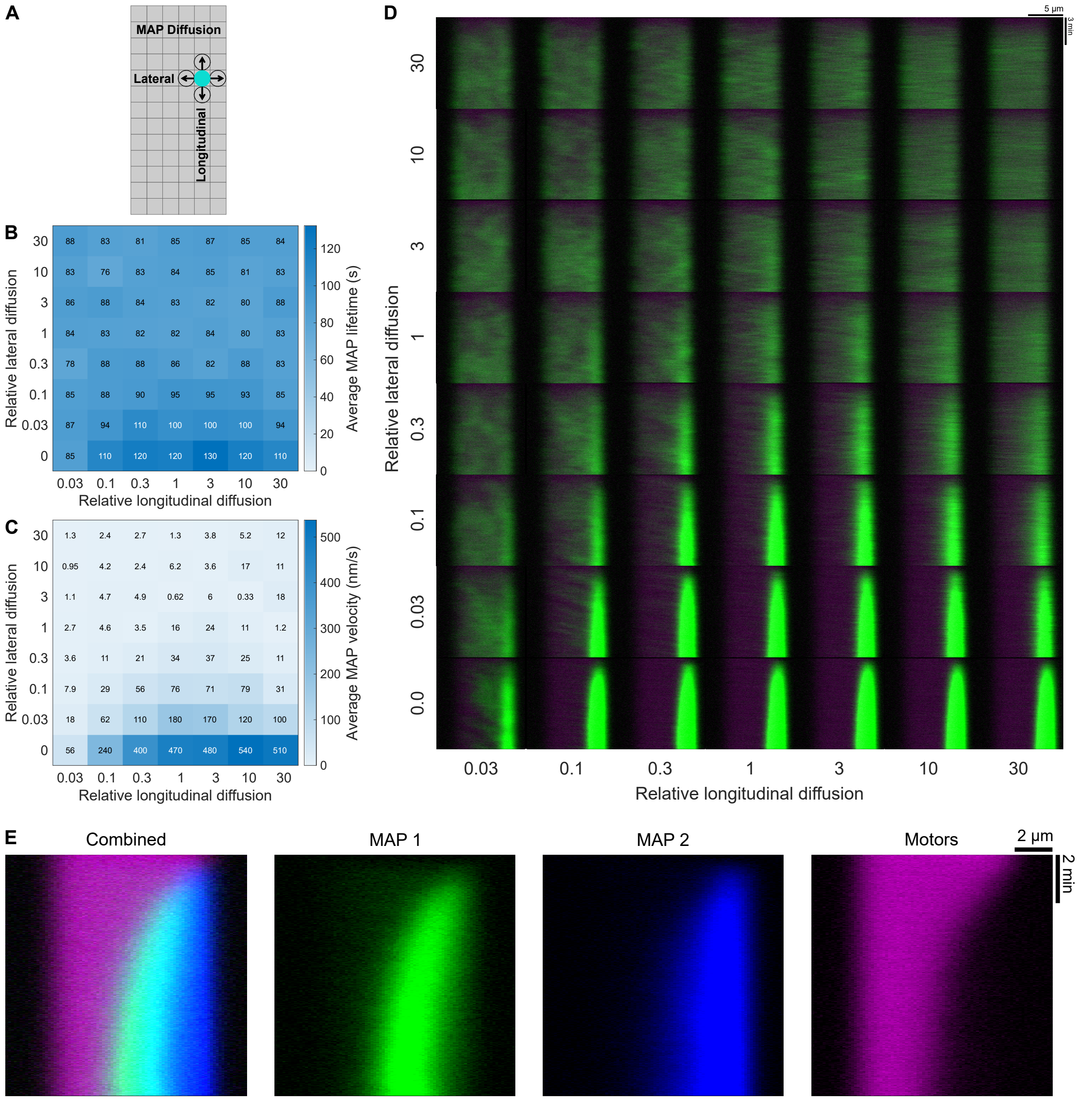


Fig. S2: **Shepherding depends on MAP diffusion and can lead to segregation of MAPs with different diffusivity.** (A) Schematic of lateral and longitudinal diffusion. (B, C) Average MAP lifetime (B) and velocity (C) as a function of relative longitudinal and lateral diffusion coefficients. (D) Simulated kymographs of MAP accumulation over time for simulations with varying longitudinal and lateral diffusion coefficients. Relative diffusion coefficients are scaled from the reference model (Table S1). MAP and motor concentration, 0.1 and 10 nM. (E) Simulated kymographs of model with two MAP species. MAP 1 (green) and MAP 2 (blue) have the parameter values of the reference model (Table S1), except that for MAP 2 the longitudinal diffusion coefficient is increased by a factor of 10 and the lateral diffusion coefficient is decreased by a factor of 10, which enhances shepherding of MAP 2. Shepherding leads to segregation of the two MAPs over time. MAP 1 and MAP 2 concentrations are 1 nM and motor concentration is 100 nM. For all simulations, microtubules are 1000 sites ≈ 8 $\mu$m long.

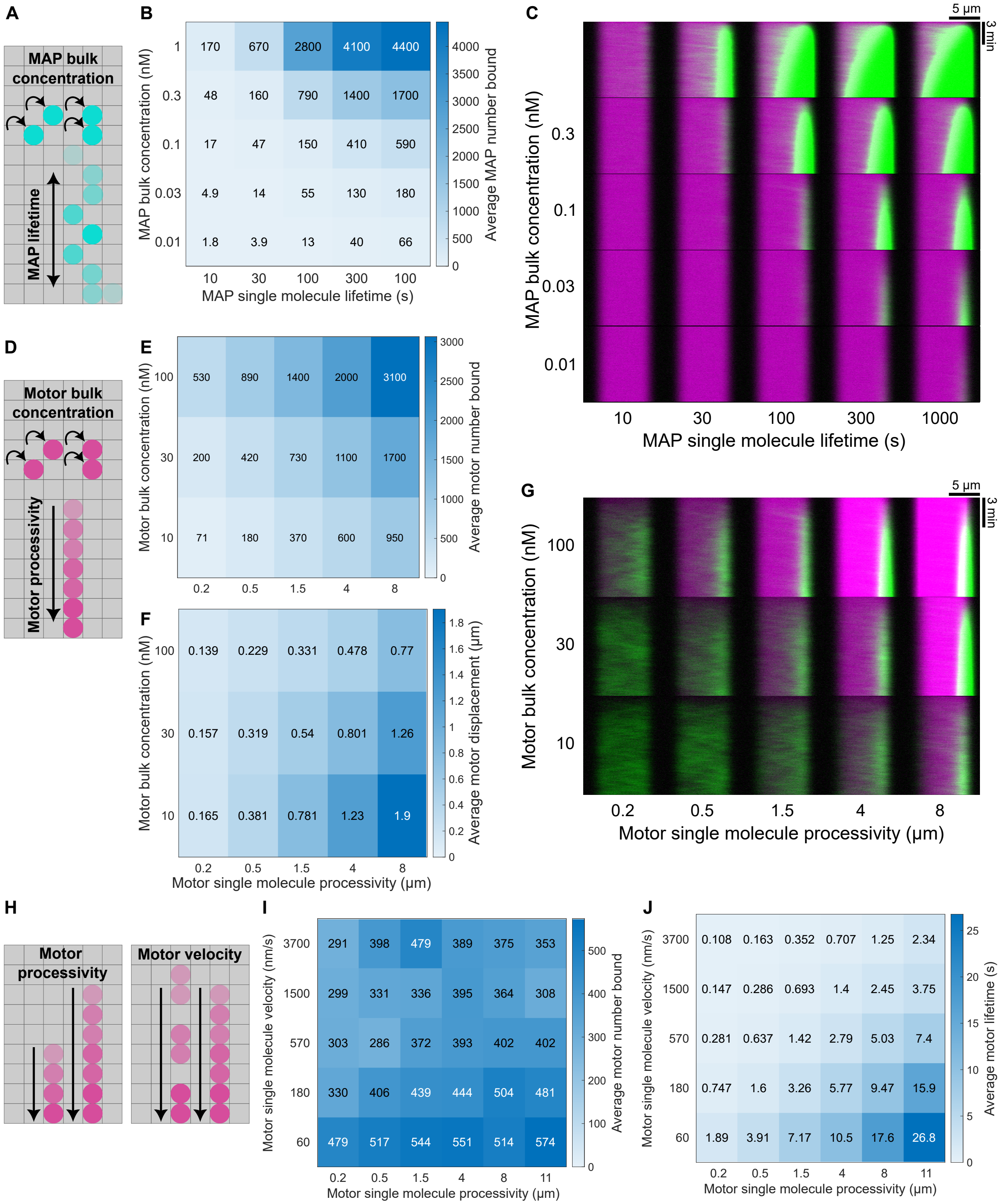
A
MAP bulk concentration
MAP lifetime
B
MAP bulk concentration (nM)
1
0.3
0.1
0.03
0.01
170 670 2800 4100 4400
48 160 790 1400 1700
17 47 150 410 590
4.9 14 55 130 180
1.8 3.9 13 40 66
10 30 100 300 100
MAP single molecule lifetime (s)
Average MAP number bound
4000 3500 3000 2500 2000 1500 1000 500 0
C
5 µm
3 min
MAP bulk concentration (nM)
0.3
0.1
0.03
0.01
10 30 100 300 1000
MAP single molecule lifetime (s)
D
Motor bulk concentration
Motor processivity
E
Motor bulk concentration (nM)
100
30
10
530 890 1400 2000 3100
200 420 730 1100 1700
71 180 370 600 950
0.2 0.5 1.5 4 8
Average motor number bound
3000 2500 2000 1500 1000 500 0
F
Motor bulk concentration (nM)
100
30
10
0.139 0.229 0.331 0.478 0.77
0.157 0.319 0.54 0.801 1.26
0.165 0.381 0.781 1.23 1.9
0.2 0.5 1.5 4 8
Motor single molecule processivity (µm)
Average motor displacement (µm)
1.8 1.6 1.4 1.2 1 0.8 0.6 0.4 0.2 0
G
5 µm
3 min
Motor bulk concentration (nM)
100
30
10
0.2 0.5 1.5 4 8
Motor single molecule processivity (µm)
H
Motor processivity
Motor velocity
I
Motor single molecule velocity (nm/s)
3700
1500
570
180
60
291 398 479 389 375 353
299 331 336 395 364 308
303 286 372 393 402 402
330 406 439 444 504 481
479 517 544 551 514 574
0.2 0.5 1.5 4 8 11
Motor single molecule processivity (µm)
Average motor number bound
500 400 300 200 100 0
J
Motor single molecule velocity (nm/s)
3700
1500
570
180
60
0.108 0.163 0.352 0.707 1.25 2.34
0.147 0.286 0.693 1.4 2.45 3.75
0.281 0.637 1.42 2.79 5.03 7.4
0.747 1.6 3.26 5.77 9.47 15.9
1.89 3.91 7.17 10.5 17.6 26.8
0.2 0.5 1.5 4 8 11
Motor single molecule processivity (µm)
Average motor lifetime (s)
25 20 15 10 5 0

Fig. S3: **Shepherding depends on the number of motors bound to microtubule, not single-motor processivity.** (A) Schematic of MAP bulk concentration and lifetime. (B) Average number of bound MAPs as a function of MAP concentration and lifetime. (C) Simulated kymographs of MAP accumulation over time for simulations with varying MAP concentration and lifetime. Motor concentration, 100 nM. (D) Schematic of motor concentration and processivity. (E, F) Average motor number bound (E) and motor displacement (F) as a function of motor concentration and lifetime. (G) Simulated kymographs of MAP accumulation over time for simulations with varying motor concentration and lifetime. MAP concentration, 0.1 nM. (H) Schematic of motor velocity and processivity. (I, J) Average motor number bound (I) and motor lifetime (J) as a function of motor concentration and lifetime. Motor number is kept approximately constant by adjusting concentration based on measured lifetime, which we note is roughly constant along each diagonal in the grid. For diagonal lines with measured lifetimes of approximately 0.1, 0.15, 0.3, 0.7, 1.5, 3, 5, 10, 15, and 30 seconds, motor concentrations are 350, 240, 120, 50, 18, 10, 5.2, 2.9, 2.0, 1.1, and 0.8 nM, respectively. For all simulations, microtubules are 1000 sites $\approx 8$ $\mu$m long.

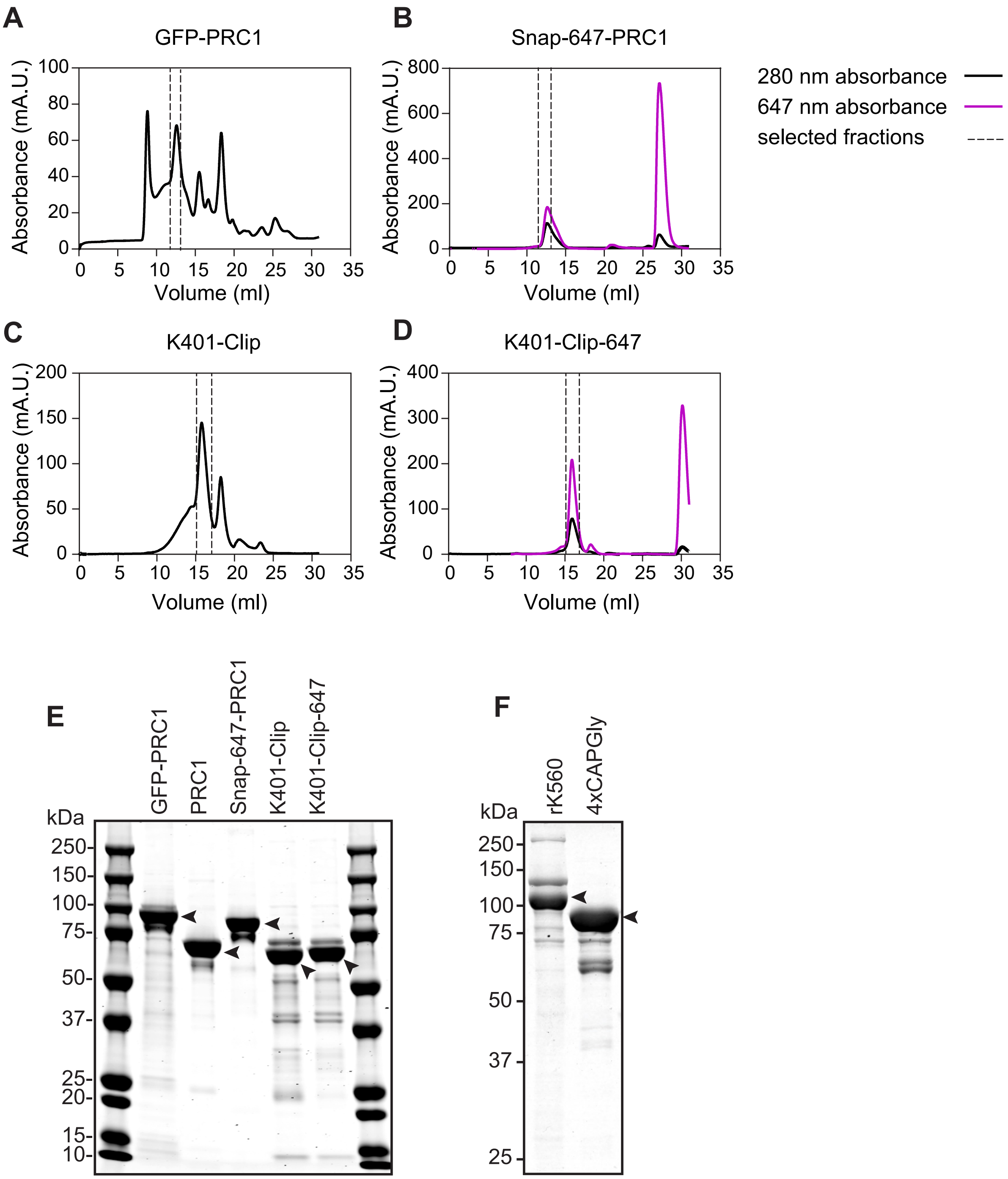


Fig. S4: **Protein purification.** (A-D) Elution profiles showing 280 nm absorbance (black) or 647 nm absorbance (magenta) from size exclusion chromatography using a Superose-6-10-300GL column for GFP-PRC1 (A), Snap-647-PRC1 (B), K401-Clip (C), and K401-Clip-647 (D). The area between the respective dashed lines indicates fractions that were pooled and concentrated. (E) Coomassie-stained PAGE gel of purified proteins: GFP-PRC1, PRC1, Snap-647-PRC1, K401-Clip, and K401-Clip-647. (F) Coomassie-stained PAGE gel of purified proteins: rKIF5C (1-560), and 4xCAPGly-GFP.

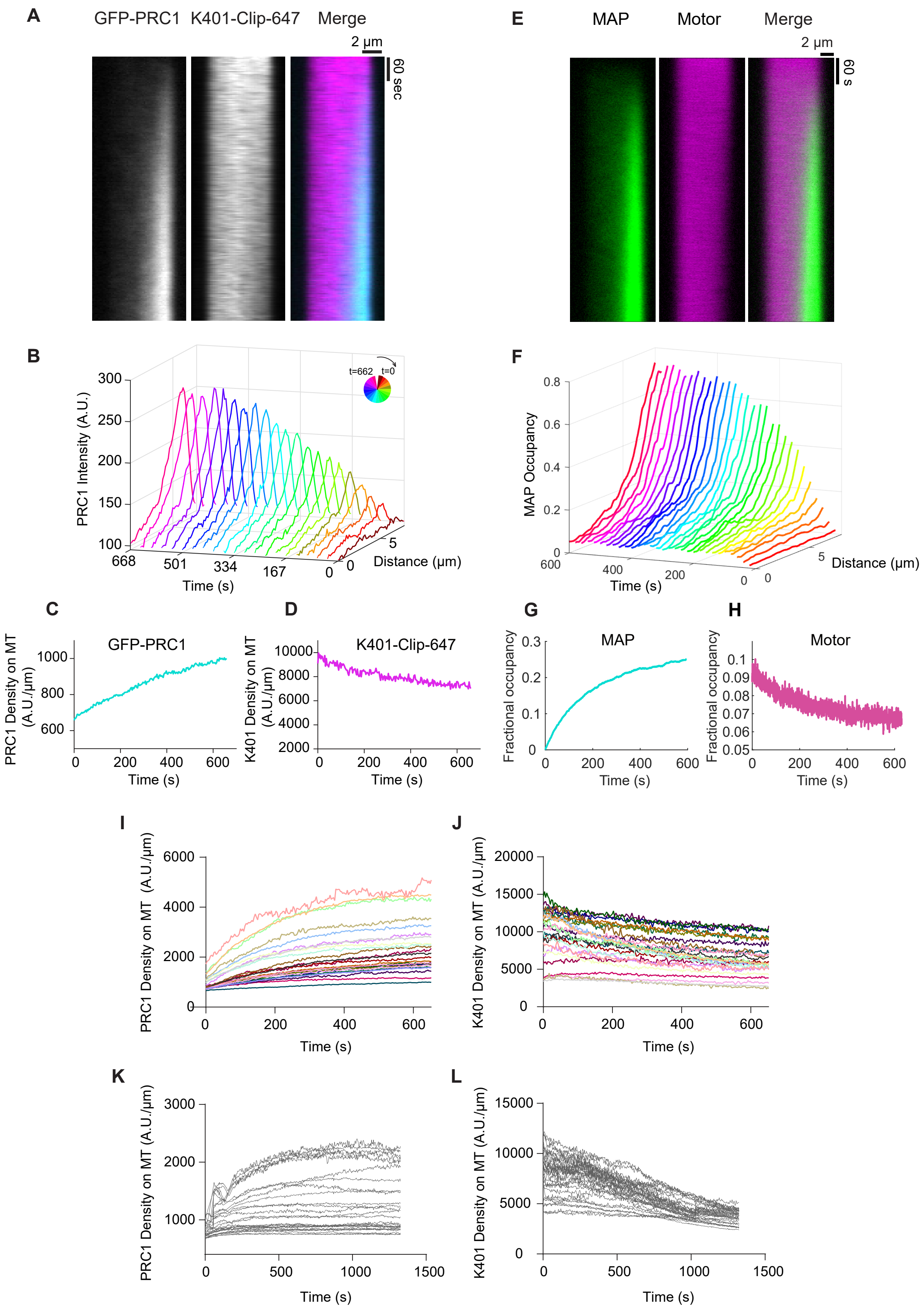
A
GFP-PRC1
K401-Clip-647
Merge
2 µm
60 sec
E
MAP
Motor
Merge
2 µm
60 s
B
PRC1 Intensity (A.U.)
t=662
t=0
Time (s)
Distance (µm)
F
MAP Occupancy
Time (s)
Distance (µm)
C
GFP-PRC1
PRC1 Density on MT (A.U./µm)
Time (s)
D
K401-Clip-647
K401 Density on MT (A.U./µm)
Time (s)
G
MAP
Fractional occupancy
Time (s)
H
Motor
Fractional occupancy
Time (s)
I
PRC1 Density on MT (A.U./µm)
Time (s)
J
K401 Density on MT (A.U./µm)
Time (s)
K
PRC1 Density on MT (A.U./µm)
Time (s)
L
K401 Density on MT (A.U./µm)
Time (s)

Fig. S5: **Dynamics of PRC1 endzone formation.** (A-D) Experimental results. (A) Representative kymograph: GFP-PRC1 (1nM, left), K401-Clip-647 (100 nM, middle) and merge (right). The horizontal and vertical scale bars are 2 $\mu$m and 60 s. (B) PRC1 intensity as a function of distance along the microtubule over time from the example in A. The time interval between curves is $\sim$30 s. The endzone forms within the first 30 s and then enriches considerably over the imaging duration of 662 seconds. (C) PRC1 density and (D) K401 density on microtubule as a function of time from the example in A. (E-H) Simulation results. MAP and motor concentrations are 0.75 nM and 30 nM. Microtubules are 1000 sites $\approx$ 8 $\mu$m long. (E) Simulated kymograph showing MAP (left), motor (middle) and merge (right). (F) MAP occupancy as a function of distance along the microtubule over time from the example in E. (G) MAP and (H) motor fractional occupancy on the microtubule as a function of time from the example in E. (I) GFP-PRC1 density on microtubules over time from experiments with 1 nM GFP-PRC1 and 100 nM K401-Clip-647. The occupancy of PRC1 increases with a characteristic time of 304 $\pm$ 130 seconds (mean $\pm$ standard deviation; N = 25, three independent experiments). (J) K401-Clip-647 density on microtubules over time from experiments with 1 nM GFP-PRC1 and 100 nM K401-Clip-647. The occupancy of K401 motors slightly decreases despite the higher concentration of K401 (N = 25, three independent experiments). (K) GFP-PRC1 density on microtubules in control experiments with 1 nM PRC1. The PRC1 occupancy increases with a characteristic time of 168 $\pm$ 84 seconds (mean $\pm$ standard deviation for N = 25, two independent experiments). (L) K401-Clip-647 density on microtubules in control experiments with 100 nM K401. The apparent density decreases over time, which is likely due to photobleaching (N = 29, two independent experiments). Control experiments were performed using identical buffer and imaging conditions.

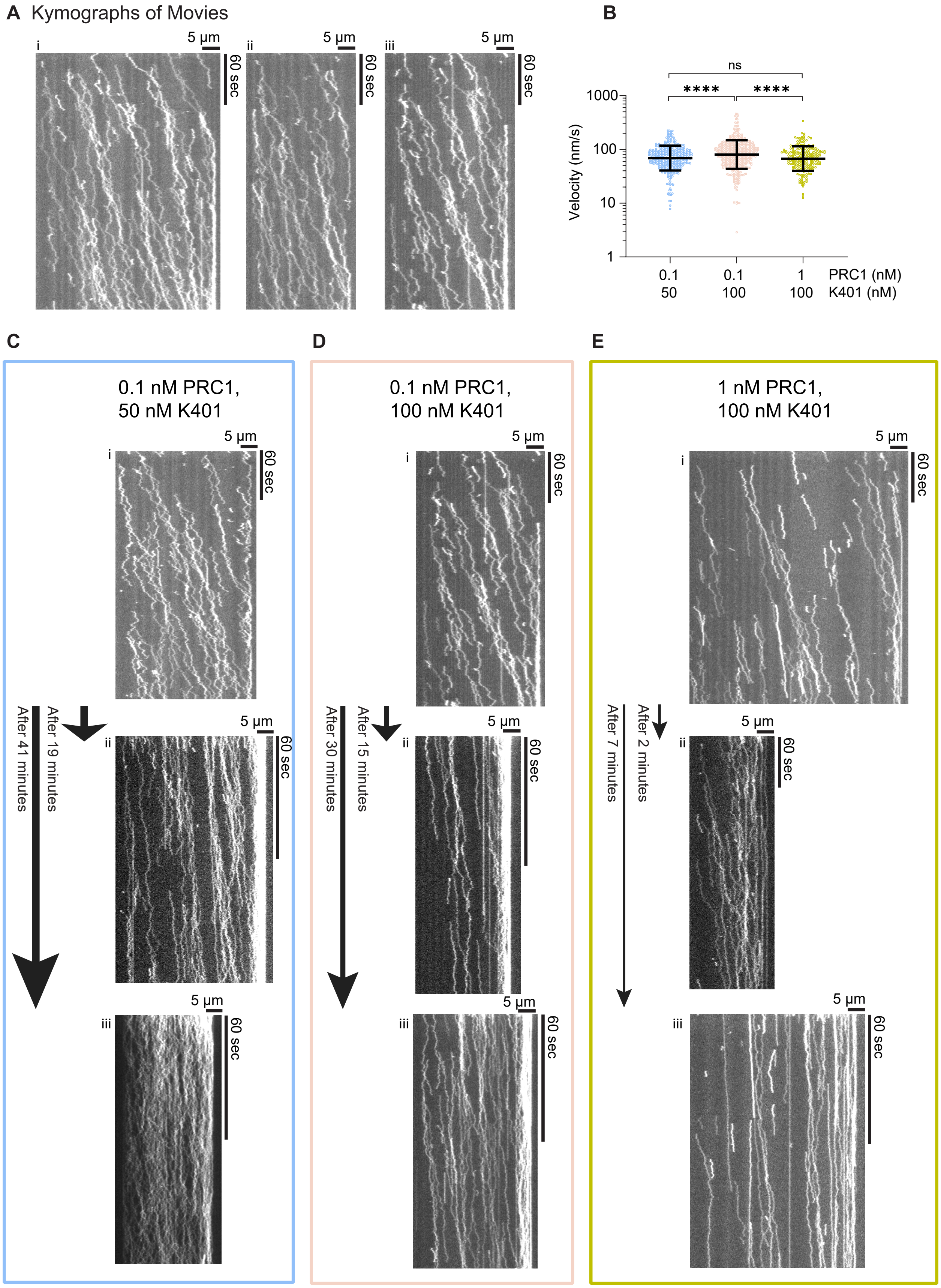
A Kymographs of Movies
i
ii
iii
5 µm
60 sec
B
ns
****
****
1000
100
10
1
Velocity (nm/s)
0.1
50
0.1
100
1
100
PRC1 (nM)
K401 (nM)
C
0.1 nM PRC1,
50 nM K401
After 19 minutes
After 41 minutes
D
0.1 nM PRC1,
100 nM K401
After 15 minutes
After 30 minutes
E
1 nM PRC1,
100 nM K401
After 2 minutes
After 7 minutes

Fig. S6: **PRC1 movement changes over time during endzone formation.**(A) Kymographs corresponding to filaments i, ii, and iii in supplementary movie 2 and movie 3, showing the biased diffusion of PRC1 in the presence of unlabeled K401 at 0.1 nM PRC1 and 100 nM K401. All kymographs are 5 minutes long and were captured immediately after protein addition to immobilized microtubules. The fraction of fluorescent PRC1 is 10%. The horizontal and vertical scale bars are 5 $\mu$m and 60 s. (B) K401 velocity as a function of K401 and PRC1 concentration for the distributions shown in Fig. 4F-H. Bars represent mean and standard deviation from fitting to a normal distribution. P-values were calculated by unpaired parametric t-test. (C-E) Representative kymographs showing biased diffusion of PRC1 at 0.1 nM PRC1 and 50 nM K401 (C), 0.1 nM PRC1 and 100 nM K401 (D), and 1 nM PRC1 and 100 nM K401 (E). (i) The top row of kymographs are 5 min long and were captured immediately after protein addition to immobilized microtubules. (ii, iii) The middle and lower rows of kymographs are 2-5 min long and were acquired at indicated times after protein addition. The arrows indicate the time interval between kymographs. The fraction of fluorescent PRC1 is 10% in A, B, and 0.5% in C. The horizontal and vertical scale bars are 5 $\mu$m and 60 s. At early time, biased diffusion of PRC1 molecules is observable that leads to endzone formation and enrichment. At intermediate time, the biased movement appears decreased. At later time, little to no PRC1 displacement was observed, indicating a stall of shepherding. The time when shepherding stalls depends on protein concentration.

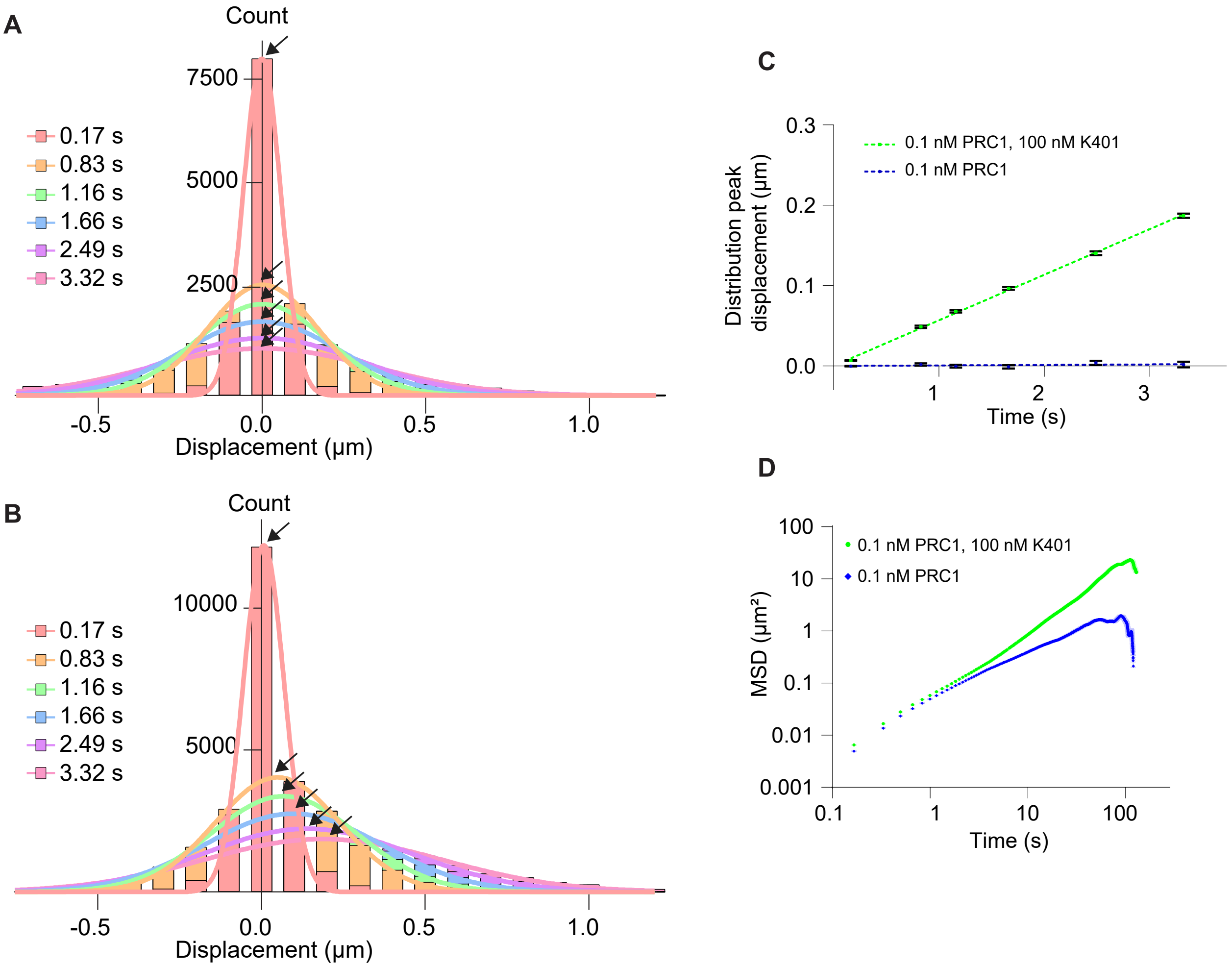


Fig. S7: **Displacement measurement of PRC1 molecules.** (A-B) Histograms of PRC1 displacement at indicated time intervals at 0.1 nM PRC1 (A) and 0.1 nM PRC1 and 100 nM K401 (B). The histograms were fitted to the normal distribution and the location of peaks is shown with arrows. (A) Mean ($\mu$m), SEM, and N for 0.17 s: -0.00038, 0.00052, 11943; 0.83 s: 0.00185, 0.00157, 11515; 1.16 s: -0.00038, 0.00188, 11301; 1.66 s: -0.00127, 0.00229, 10980; 2.49 s: 0.00368, 0.00288, 10445; 3.32 s: 0.001816, 0.00342, 9910. N=136 tracks, bin size 0.1 $\mu$m. (B) Mean ($\mu$m), SEM, and N for 0.17 s: 0.006863, 0.000435, 20316; 0.83 s: 0.04876, 0.001352, 19728; 1.16 s: 0.06807, 0.00161, 19434; 1.66 s: 0.09648, 0.001956, 18993; 2.49 s: 0.14030, 0.00236, 18258; 3.32 s: 0.18690, 0.00276, 17523. N=157 tracks, bin size 0.1 $\mu$m. (C) The mean displacement of PRC1 molecules as a function of time for 0.1 nM PRC1 (blue), and 0.1 nM PRC1 and 100 nM K401 (green) for 0.17, 0.83, 1.16, 1.66, 2.49, and 3.32 s intervals displayed in A-B. Linear fitting gives corresponding slope ($\mu$m/s) and 95% CI of 0.00063 [-0.00031, 0.0016] and 0.05676 [0.05598, 0.05755], respectively. Error bars show SEM. (D) Mean squared displacement (MSD) of PRC1 molecules as a function of time are shown for 0.1 nM PRC1 (blue), and 0.1 nM PRC1 and 100 nM K401 (green). N=136 and 157 tracks, respectively. Error bars show SEM.

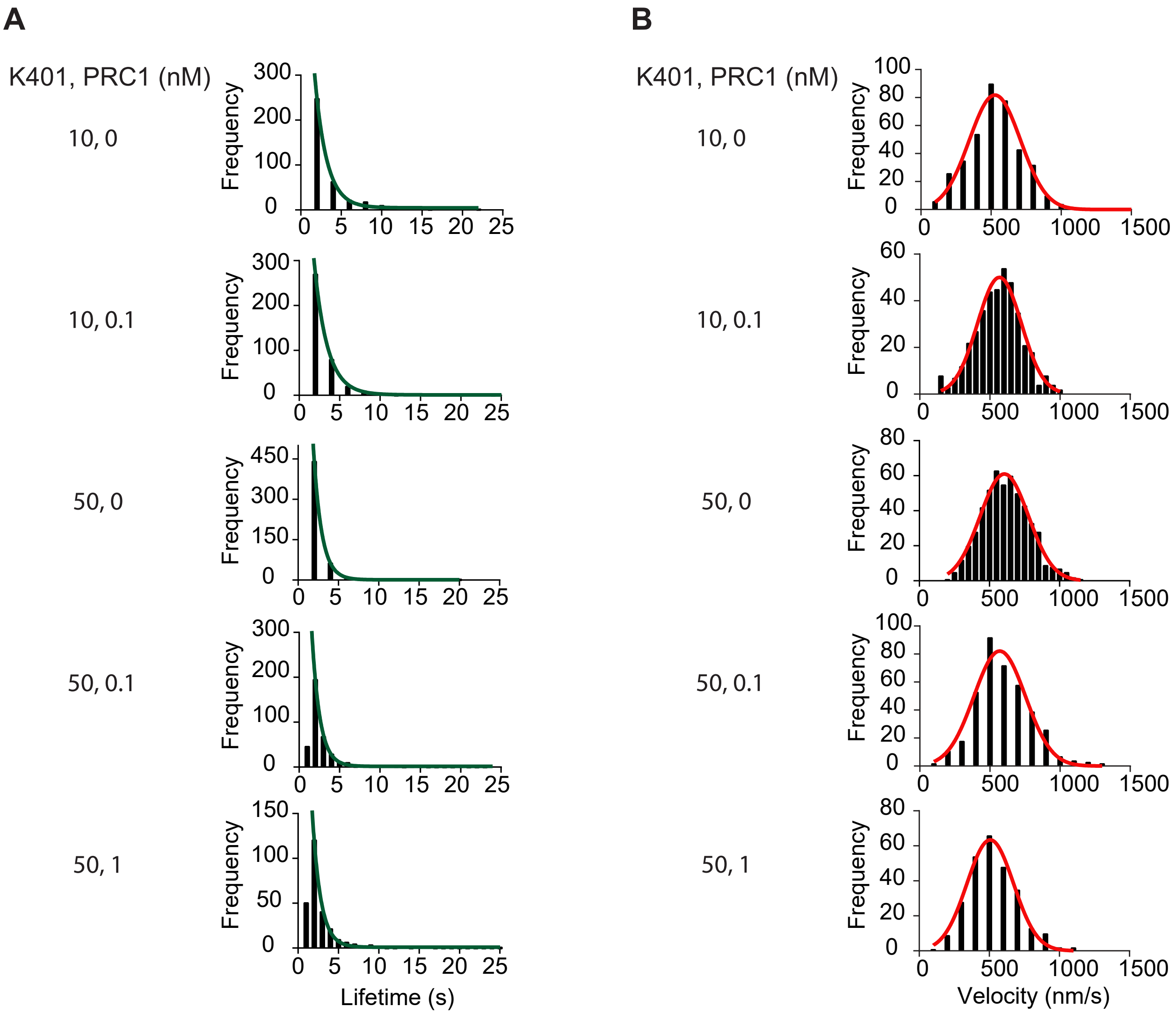


Fig. S8: **Histograms of K401 lifetime and velocity for data shown in Fig. 5.**(A) Histograms of K401 lifetime at indicated concentrations of K401 and PRC1. The histograms were fitted to an exponential distribution (green). Mean, 95% confidence interval range, N, and bin sizes are: 10 nM K401: 1.473 [1.27, 1.69] s, N=380, 2; 10 nM K401 and 0.1 nM PRC1: 1.63 [1.58, 1.69] s, N=398, 2; 50 nM K401: 1.02 s [0.97, 1.08] s, N=524, 2; 50 nM K401 and 0.1 nM PRC1: 0.99 [0.94, 1.05] s, N=387, 1; 50 nM K401 and 1 nM PRC1: 1.04 [0.96, 1.13] s, N=268, 1. N is the number of tracks analyzed in three independent experiments in each condition. (B) Histograms of K401 velocity at indicated concentrations of K401 and PRC1. The histograms were fitted to a normal distribution (red). Mean, standard deviation, N values, and bin sizes are: 10 nM K401: 527 ± 183 nm/s, 380, 100; 10 nM K401 and 0.1 nM PRC1: 568 ± 156 nm/s, N=398, 50; 50 nM K401: 606 ± 172 nm/s, N=524, 50; 50 nM K401 and 0.1 nM PRC1: 570 ± 184 nm/s, N=387, 100; 50 nM K401 and 1nM PRC1: 507 ± 166 nm/s, N=268, 100. N is the number of tracks analyzed in three independent experiments in each condition. The K401 velocity does not vary significantly between these conditions.

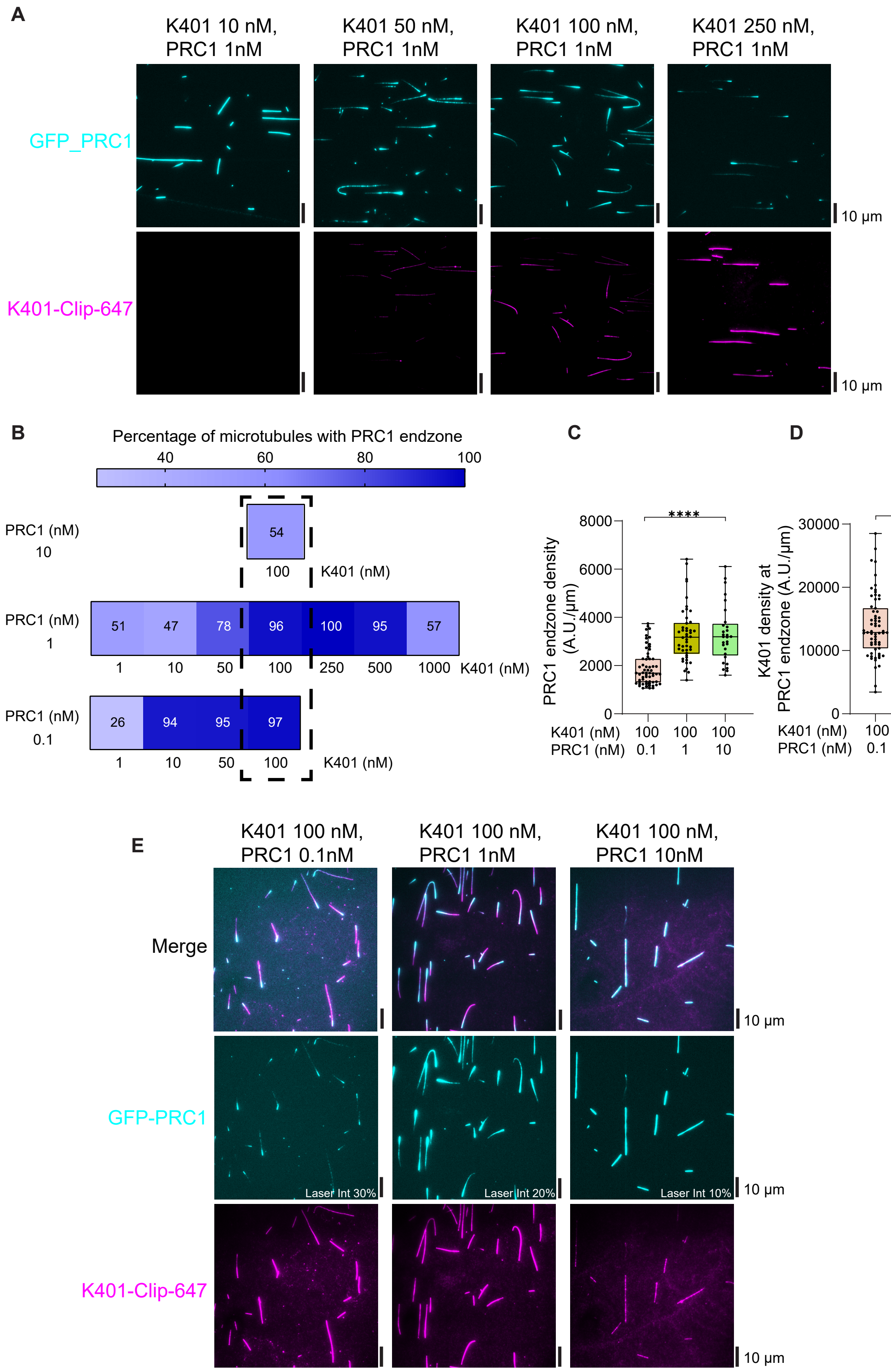
A
K401 10 nM, PRC1 1nM
K401 50 nM, PRC1 1nM
K401 100 nM, PRC1 1nM
K401 250 nM, PRC1 1nM
GFP_PRC1
K401-Clip-647
10 μm
10 μm
B
Percentage of microtubules with PRC1 endzone
40
60
80
100
PRC1 (nM) 10
54
100
K401 (nM)
PRC1 (nM) 1
51
47
78
96
100
95
57
1
10
50
100
250
500
1000
K401 (nM)
PRC1 (nM) 0.1
26
94
95
97
1
10
50
100
K401 (nM)
C
PRC1 endzone density (A.U./μm)
8000
6000
4000
2000
0
****
K401 (nM) 100 100 100
PRC1 (nM) 0.1 1 10
D
K401 density at PRC1 endzone (A.U./μm)
30000
20000
10000
0
****
K401 (nM) 100 100 100
PRC1 (nM) 0.1 1 10
E
K401 100 nM, PRC1 0.1nM
K401 100 nM, PRC1 1nM
K401 100 nM, PRC1 10nM
Merge
GFP-PRC1
K401-Clip-647
Laser Int 30%
Laser Int 20%
Laser Int 10%
10 μm
10 μm
10 μm

Fig. S9: **Additional images and analysis, for data shown in Fig. 6.** (A) Representative fields of view in Fig. 6A shown with the same fluorescence intensity scaling for each image GFP-PRC1 in cyan, K401-Clip-647 in magenta. GFP-PRC1 fluorescence intensity on microtubules decreases with increasing K401-Clip-647 concentration, while K401 fluorescence intensity increases. Scale bars, 10 $\mu$m. (B) The percentage of microtubules with a visible PRC1 endzone at steady state from Fig. 6B with additional data for 10 nM PRC1 and 100 nM K401 (N = 61). The data are from three independent experiments. (C-D) Steady state endzone density of GFP-PRC1 (C) and K401-Clip-647 density in the PRC1 endzone (D) for 100 nM K401-Clip-647 with 0.1, 1, and 10 nM GFP-PRC1. Box edges, middle line, and whiskers are 25th and 75th percentiles, median and minimum and maximum values, respectively. P-values were calculated by unpaired nonparametric Kolmogorov-Smirnov t-test. GFP-PRC1 endzone density mean, standard deviation, and N are 100 nM K401, 0.1 nM PRC1: 1911 $\pm$ 736 A.U./$\mu$m, N=61; 100 nM K401, 1 nM PRC1: 3279 $\pm$ 1134 A.U./$\mu$m, N=46; and 100 nM K401, 10 nM PRC1: 3304 $\pm$ 1180 A.U./$\mu$m, N=29. The data are from three independent experiments. K401 density mean, standard deviation, and N are 100 nM K401, 0.1 nM PRC1: 13862 $\pm$ 4971 A.U./$\mu$m, N=63; 100 nM K401, 1 nM PRC1: 8891 $\pm$ 3809 A.U./$\mu$m, N=46; and 100 nM K401, 10 nM PRC1: 5881 $\pm$ 1695 A.U./$\mu$m, N=29. The data are from three independent experiments. (E) Representative fields of view at steady state for assays with 100 nM K401-Clip-647 and varying GFP-PRC1. The fluorescence intensity scaling is the same for each image, but data were acquired at varying 488 nm laser intensity excitation of GFP-PRC1. 488 nm laser intensity percentages are 100 nM K401, 0.1 nM PRC1: 30%; 100 nM K401, 1 nM PRC1: 20%; 100 nM K401, 10 nM PRC1: 10%, as labeled in middle row. K401-Clip-647 fluorescence intensity decreases with increasing PRC1 concentration. GFP-PRC1 fluorescence intensity increases with its concentration as expected. Scale bars, 10 $\mu$m.

#### Movie captions

Movie S1: Gliding assay shows accumulated GFP-PRC1 at the rear (plus end) of microtubules moved by surface immobilized K401 motors. GFP-PRC1 and K401-Clip-647 are displayed in cyan and magenta, respectively. The scale bar is 10 $\mu$m. The time interval is $\sim$6 sec, and the total duration of the movie is $\sim$10 min.

Movie S2: Biased diffusion of PRC1 molecules along immobilized microtubules at 0.1 nM PRC1 and 100 nM K401. The fraction of fluorescent Snap-647-PRC1 (gray) is 10%, while K401 motors are non-fluorescent. The scale bars are 5 $\mu$m. The time interval is $\sim$160 msec, and the total duration of the movie is $\sim$5 min.

Movie S3: Biased diffusion of PRC1 molecules along immobilized microtubules at 0.1 nM PRC1 and 100 nM K401. The fraction of fluorescent Snap-647-PRC1 (gray) is 10%, while K401 motors are non-fluorescent. The scale bars are 5 $\mu$m. The time interval is $\sim$160 msec, and the total duration of the movie is $\sim$5 min.

[22] Nandini Mani, Michelle F. Marchan, and Radhika Subramanian. Simultaneous Visualization of the Dynamics of Crosslinked and Single Microtubules In Vitro by TIRF Microscopy. *Journal of Visualized Experiments (JoVE)*, (180):e63377, February 2022.

[23] Stefan Niekamp, Jongmin Sung, Walter Huynh, Gira Bhabha, Ronald D. Vale, and Nico Stuurman. Nanometer-accuracy distance measurements between fluorophores at the single-molecule level. *Proceedings of the National Academy of Sciences*, 116(10):4275–4284, March 2019.

[24] Stefan Niekamp, Nico Stuurman, Nan Zhang, and Ronald D. Vale. Three-color single-molecule imaging reveals conformational dynamics of dynein undergoing motility. *Proceedings of the National Academy of Sciences*, 118(31):e2101391118, August 2021.
